# Whole-Embryo 3D Quantification Reveals Flexible Cellular Scaling and Conserved Tissue Architecture in *Xenopus* Species

**DOI:** 10.64898/2026.06.16.732511

**Authors:** Hugo Mauricio, Mariia Diakova, Max Brambach, Cora Anderson, Kseniya Petrova, Cecilia A. De Araujo, Iva Simeonova, Geneviève Almouzni, Leonid Peshkin, Jose G. Abreu

## Abstract

How embryos with markedly different absolute sizes and cell numbers establish comparable tissue organization during development remains a fundamental question in developmental biology. To address this question, we compared two closely related *Xenopus* species that differ substantially in embryonic size, *Xenopus laevis* and *Xenopus tropicalis*. We generated a whole-embryo quantitative 3D atlas of cell allocation, spatial organization, and mitotic dynamics at key time points between gastrulation to tailbud stages. Using tissue clearing, and 3D imaging we tracked single-nucleus coordinates across developmental milestones to resolve how body plans adapt to organismal scale. We show that embryonic scaling is not achieved through simple proportional changes in cell number. Instead, the smaller *X. tropicalis* embryo is characterized by a distinct high-density tissue organization associated with a persistently higher mitotic index (∼1.4-fold higher than in *X. laevis* at both gastrula and tailbud stages). Across development, this is accompanied by a near-doubling of cell number in *X. tropicalis* without a proportional increase in embryo volume. We quantify tissue organization and find species-specific cellular architectures during gastrulation that largely converge by the tailbud stage. At this stage, homologous tissues display broadly similar structural profiles despite persistent differences in embryo size, cell number, and density. Together, our findings reveal that closely related vertebrate embryos can follow distinct cellular organization trajectories while converging toward comparable tissue architecture, providing a quantitative framework for understanding robust body plan formation across divergent physical scales.

**Key Findings:**

- *The number of cells in X. tropicalis increase*s by nearly double between the gastrula and tailbud stages, even though the volume of the embryo does not increase accordingly.
- *X. laevis has a mitotic index that is about 1.4 times lower than that of X. tropicalis* at both the gastrula and tailbud stages.
- Systematic differences in tissue density are observed among the germ layers within the embryo (consistently in both species), with the neuroectoderm being among the most densely packed and the endoderm among the least dense.
- Cell number, density, and structure vary substantially among germ layers during gastrulation; however, these differences between germ layers diminish as development progresses.
- In the tailbud stage the homologous tissues in *X. laevis* and *X. tropicalis* show generally similar structural arrangements even though there are continuous differences in embryo size and cell number.

## INTRODUCTION

One of the most fundamental questions in developmental biology is how embryos of different species and different sizes reliably produce almost identical body plans. This phenomenon, called developmental scaling, implies the existence of robust mechanisms that coordinate cell number, tissue size, and spatial organization across species and across individuals within a species (Wolpert, 1969; Čapek & Müller, 2019; Timoshina & Zaraisky, 2026). Specifically, recent work has shown that accurate scaling of morphogen gradients requires dedicated “scaler” genes whose expression is tuned to embryo size. Several scaler genes have now been identified in *Xenopus* and sea urchin, revealing the mechanisms by which they rescale e.g. Chordin/BMP gradients with embryo size (Orlov et al., 2022; Timoshina et al., 2025; Timoshina & Zaraisky, 2026). During the transition from gastrulation through neurulation to the tailbud stage, the three primary germ layers, ectoderm, mesoderm, and endoderm, are progressively specified, allocated, and patterned to architect the vertebrate body axis. Whether these processes scale linearly with cell number or rely on conserved spatial organization “design rules” (Mayr et al., 2019) that are independent of absolute embryo size remains poorly understood.

Classic experimental studies demonstrated that embryonic form can remain remarkably robust despite major alterations in cell size and cell number. In pioneering work, Fankhauser showed that polyploid amphibian embryos develop with fewer but substantially larger cells while maintaining nearly normal tissue and organ dimensions (Fankhauser, 1945; Fankhauser, 1972). Fankhauser’s experiments on newt embryos showed that making cells larger by increasing ploidy did not lead to a corresponding increase in the overall size of the animal. Instead, tissues and organs such as the pronephric duct retained their normal dimensions, even though the number of cells forming the duct dropped dramatically—in pentaploid embryos, as few as one to three cells had to enclose a lumen that normally required three to eight cells. More recent studies in vertebrate and invertebrate systems have further shown that morphogen gradients, tissue proportions, and developmental dynamics can adapt to changes in embryo size through scale-invariant regulatory and mechanical mechanisms (Ben-Zvi et al., 2011; Almuedo-Castillo et al., 2018; Heisenberg & Bellaïche, 2013; Orlov et al., 2022; Timoshina et al., 2025). Together, these findings suggest that embryonic development may rely on conserved organizational principles that preserve tissue architecture independently of absolute embryo size or cell number. Recent advances in whole-embryo imaging and computational analysis have enabled reconstruction of vertebrate development at cellular resolution, revealing coordinated cell behaviors and tissue dynamics across entire embryos (Papan et al., 2007; McDole et al., 2018; Mayr et al., 2019). However, quantitative comparisons that integrate absolute cell number, tissue allocation, local cellular organization, and proliferative activity across naturally different embryonic scales remain limited.

The genus *Xenopus* provides an exceptional comparative framework for investigating how differences in embryonic size and cell number are accommodated while preserving overall tissue organization and developmental form. *Xenopus laevis*, a pseudotetraploid with large (∼1.2 mm diameter) yolk-rich eggs, and *Xenopus tropicalis*, a diploid with smaller (∼0.7 mm) eggs, exhibit substantial differences in egg size, genome content, cell volume, and total embryonic volume (Levy & Heald, 2010; Gibeaux et al., 2018). Comparative studies across *Xenopus* species further indicate that size relationships change during embryogenesis, with distinct scaling regimes emerging as development progresses (Miller et al., 2023). Despite these differences, the two species ultimately establish broadly comparable body plans, although their developmental trajectories differ in several quantitative aspects. Their close evolutionary relationship, combined with marked differences in embryonic scale, therefore provides a natural comparative system for investigating how robust tissue organization emerges across distinct cellular and developmental trajectories.

Whole-embryo, single-cell-resolution quantification has previously been achieved in several other model systems, each exploiting features favorable to imaging or lineage tracking: automated lineage tracing in the transparent, invariant embryo of C. elegans (Bao et al., 2006); a registered whole-embryo gene-expression point-cloud atlas in the Drosophila blastoderm (Fowlkes et al., 2008); light-sheet-based nuclear tracking through gastrulation in zebrafish (Keller et al., 2008) and post-implantation mouse development (McDole et al., 2018); and whole-embryo cell-shape segmentation revealing that ascidian embryos maintain invariant tissue architecture via conserved cell-cell contact rules despite substantial individual variation in absolute cell geometry (Guignard et al., 2020). Xenopus, owing to its yolk-laden opacity, has largely been excluded from this line of whole-organism quantification until recently.

Early quantitative studies in *Xenopus laevis* established that cell number is tightly regulated during gastrulation and neurulation (Cooke, 1979a, 1979b), but these analyses relied on manual counting in serial sections, limiting throughput and spatial resolution. More recently, advances in tissue clearing—such as the iDISCO protocol—combined with light-sheet microscopy and deep-learning segmentation tools like StarDist3D have finally bridged this gap, enabling whole-embryo, single-nucleus 3D quantification in opaque systems (Renier et al., 2014; Schmidt et al., 2018).

In this study, we established an optimized pipeline combining hydrogen peroxide bleaching, far-red nuclear staining with TO-PRO-3, and ethyl cinnamate-based clearing (iDISCO) to perform high-resolution in toto 3D confocal imaging across *Xenopus* embryogenesis. By pairing this volumetric imaging with deep-learning instance segmentation via StarDist3D, we quantified absolute nuclei counts, tissue-specific cell allocation, 3D Voronoi-derived packing metrics, and anaphase-like mitotic events across developmental stages—stages 11.5, 15, and 23 in *Xenopus laevis*, and stages 11.5 and 23 in *Xenopus tropicalis*. We set out to answer three main questions: (1) does germ-layer cell allocation scale proportionally with absolute embryo volume across species? (2) how do local spatial organization properties, such as spatial order, cell packing, and geometry, evolve across developmental stages and physical scales? (3) what cellular strategies enable species with vastly different egg sizes and cell numbers to converge upon equivalent body plans?

Our quantitative analysis reveals that embryonic scaling in *Xenopus* is not driven by simple linear cell multiplication or rigid, invariant density. Instead, we show that *X. tropicalis* enter morphogenesis with distinct cellular architectures, tissue allocations, and packing states. These differences are progressively remodeled during development: by the early tailbud stage, homologous tissues display broadly similar architectural profiles despite persistent differences in absolute embryo size, cell number, and cellular density. Together, our findings suggest that developmental scaling does not require identical initial cellular conditions, but instead involves the progressive convergence of distinct cellular trajectories toward a conserved tissue-level organization. This quantitative framework helps resolve key questions arising from previous transcriptional studies of early *Xenopus* development (Yanai et al., 2011; Petrova et al., 2024). Specifically, by establishing absolute cell counts and spatial distributions, it provides a physical reference to evaluate lineage sampling in single-cell datasets and accounts for populations underrepresented due to mitotic silencing.

## RESULTS

### 1. 3D reconstruction and quantitative comparative analysis of cell number and tissue volume

To quantitatively map *Xenopus* embryonic architecture despite tissue opacity and autofluorescence, we established an integrated pipeline combining hydrogen peroxide bleaching, TO-PRO-3 nuclear staining, iDISCO-based ethyl cinnamate clearing, multi-view spinning disk microscopy (Mazière et al., 1996; Renier et al., 2014; Masselink and Tanaka, 2023), and deep-learning segmentation via StarDist3D (Schmidt et al., 2018; Weigert et al., 2020). Overcoming the limitations of traditional manual counts (Cooke, 1979a), this highly efficient workflow achieved a validated segmentation F1-score of 98.17% (Figure S1 A–C), converting non-quantitative 3D raw signal reconstructions into a fully segmented, quantitative point-cloud model (Figure 1A). These spatial coordinates were then intersected with manually annotated germ layers, quantitative 3D embryonic models across developmental stages—stages 11.5, 15, and 23 in *Xenopus laevis*, and stages 11.5 and 23 in *Xenopus tropicalis* (Figure 1B). To facilitate open-access exploration of our 3D dataset, we developed a WebGL-based online visualization platform. The platform includes an interactive viewer (https://xenopus.hms.harvard.edu/viewer.html) dedicated to germ-layer and lineage-annotated point clouds. The interactive portal enables continuous 360-degree rotation, multi-axis slicing, real-time opacity adjustment, and dynamic zoom control. Crucially, the web rendering engine supports virtual camera fly-through navigation, allowing users to enter internal embryonic volumes—such as the deep endodermal core or the inner neuroectoderm layers—and observe single-nucleus spatial arrangements, mitotic pairs, and local Voronoi architectures in situ at cellular resolution without performance degradation.

**Figure 1.**
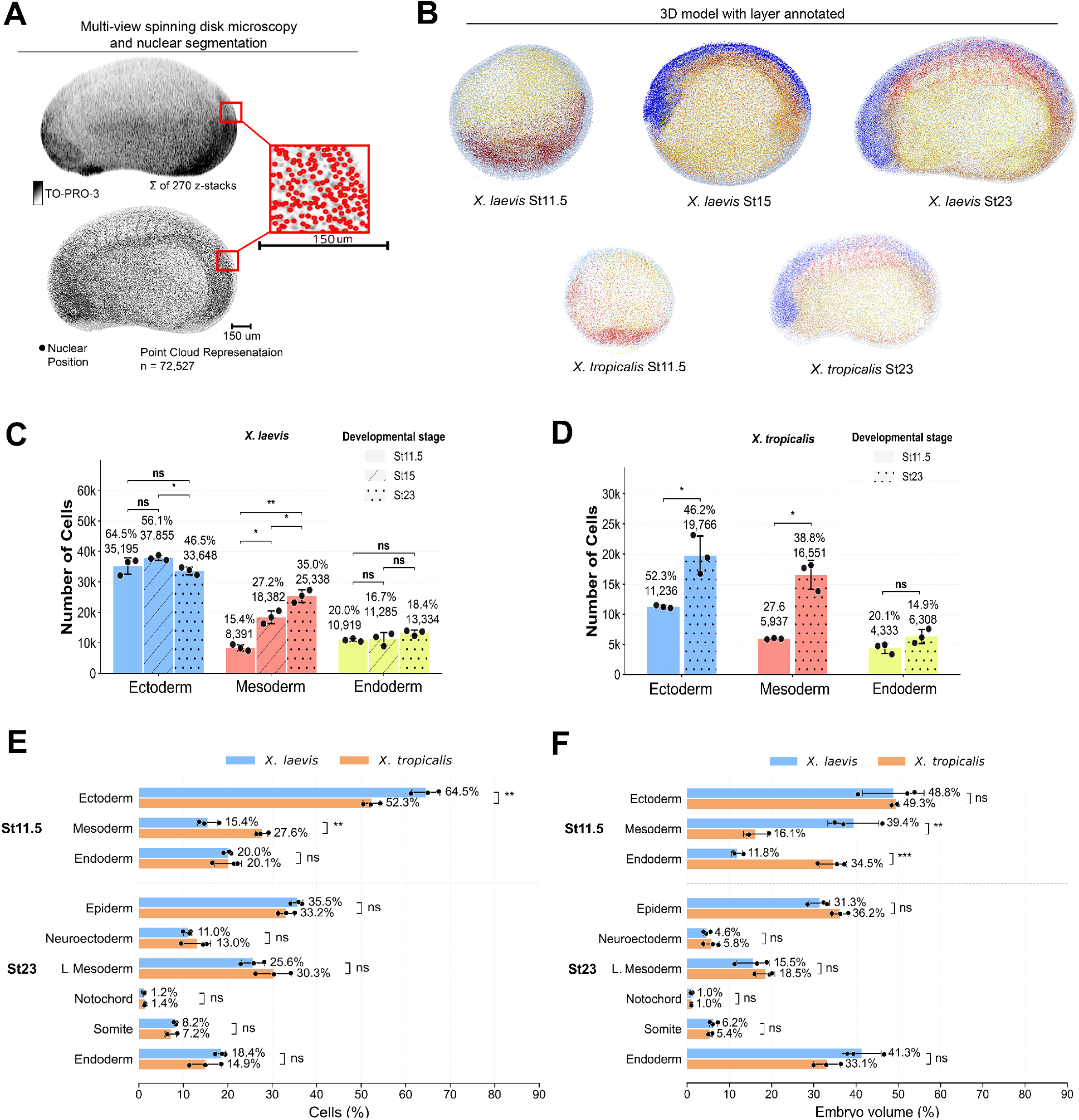
Quantitative 3D nuclear mapping reveals conserved germ-layer volume fractions despite species-specific cellular scales in *Xenopus* development. **(A)** High-resolution imaging and nuclear segmentation pipeline showing representative maximum intensity projections (sum of z-stacks) of TO-PRO-3 far-red nuclear staining in cleared *Xenopus* embryos, illustrating the conversion from raw confocal images to isolated nuclear centroids and 3D point-cloud representations (n = 72,527 centroids). **(B)** Reconstructed 3D point-cloud models color-coded by germ layer and derived lineages across embryonic stages in *Xenopus laevis* (Stage 11.5 gastrula, Stage 15 neurula, and Stage 23 tailbud) and *Xenopus tropicalis* (Stage 11.5 and Stage 23). An interactive 3D WebGL viewer for exploring these germ-layer annotated models across stages and species, including real-time virtual camera fly-through inspection, is publicly available at https://xenopus.hms.harvard.edu/viewer.html. **(C–D)** Absolute cell counts in Ectoderm, Mesoderm, and Endoderm across developmental stages for (C) *X. laevis* and (D) *X. tropicalis*, displaying raw nuclear numbers (n = 3 embryos per stage; mean ± SD). Note that at Stage 23 (St23), values for broad germ-layer categories represent the combined sum of their derived sub-lineages shown in **E** and **F** (e.g., in *X. laevis*, total Mesoderm at St23 corresponds to Lateral Mesoderm [25.6%] + Notochord [1.2%] + Somite [8.2%] = 35.0%). Note that at Stage 23 (St23), values for broad germ-layer categories represent the combined sum of their derived sub-lineages shown in **E** and **F** (e.g., in *X. laevis*, total Mesoderm at St23 corresponds to Lateral Mesoderm [25.6%] + Notochord [1.2%] + Somite [8.2%] = 35.0%). **(E)** Relative cellular proportions (%) allocated to primary germ layers at Stage 11.5 and detailed sub-lineages at Stage 23 compared between *X. laevis* (blue) and *X. tropicalis* (orange). **(F)** Relative embryonic volume fractions (%) occupied by germ layers and derived tissue structures compared between species across morphogenesis. Data are presented as mean ± SD (n = 3); statistical significance evaluated by Student’s t-test (ns = not significant,*p < 0.05, **p < 0.01, ***p < 0.001).

Detailed lineage tracking of raw cell counts in *X. laevis* highlights the precise tissue partitioning underlying these developmental transitions (Figure S2 A-C). At gastrulation (st 11.5), the embryo is predominantly composed of ectodermal cells (35,195 ± 2,666 cells), followed by endoderm (10,919 ± 496 cells) and mesoderm (8,391 ± 1,034 cells) (Figure S2A). By neurulation (st 15), spatial sub-classification revealed 24,481 ± 1,522 epidermal and 13,374 ± 1,252 neuro-ectodermal cells, while the mesoderm compartmentalized into lateral mesoderm (15,093 ± 1,364 cells), somites (2,294 ± 935 cells), and notochord (995 ± 194 cells) alongside 11,285 ± 2,107 endodermal cells (Figure S2B). By stage 23, the mesodermal lineage experienced significant expansion, driven primarily by lateral mesoderm (18,561 ± 2,076 cells) and somites (5,939 ± 154 cells), alongside 25,701 ± 694 epidermal, 7,946 ± 576 neuro-ectodermal, 838 ± 164 notochord, and 13,334 ± 886 endodermal cells (Figure S2C).

Similarly, tissue-specific lineage quantification in *X. tropicalis* demonstrated a distinct cellular trajectory toward the tailbud stage (Figure S2D-E). At stage gastrula 11.5, raw counts comprised 11,236 ± 239 ectodermal, 5,937 ± 114 mesodermal, and 4,333 ± 849 endodermal cells (Figure S2D). By tailbud stage 23, spatial mapping resolved 14,169 ± 1,750 epidermal, 5,597 ± 1,581 neuro-ectodermal, 12,909 ± 1,913 lateral mesodermal, 3,056 ± 473 somitic, 587 ± 86 notochordal, and 6,308 ± 1,174 endodermal cells (Figure S2E).

Building upon these tissue-level dynamics, cross-species comparison of consolidated cell counts and tissue volumes reveals key evolutionary and scaling principles (Figure 1C–F; n = 3, mean ± SD). *X. laevis* maintains a total cell population roughly double that of *X. tropicalis* throughout development (Figure 1C–D). At stage 11.5, *X. laevis* transitions from 35,195 ± 2,666 (64.52% ± 3.14%) ectodermal, 8,391 ± 1,034 (15.44% ± 2.34%) mesodermal, and 10,919 ± 496 (20.04% ± 0.90%) endodermal cells to 33,648 ± 1,252 (46.54% ± 2.23%) ectodermal, 25,338 ± 2,058 (35.02% ± 2.53%) mesodermal, and 13,334 ± 886 (18.43% ± 1.14%) endodermal cells at stage 23 (Figure 1C). In contrast, *X. tropicalis* starts at 11,236 ± 239 (52.30% ± 1.92%) ectodermal, 5,937 ± 114 (27.65% ± 1.39%) mesodermal, and 4,333 ± 849 (20.05% ± 3.06%) endodermal cells, reaching 19,766 ± 3,227 (46.21% ± 4.89%) ectodermal, 16,551 ± 2,376 (38.84% ± 5.19%) mesodermal, and 6,308 ± 1,174 (14.95% ± 3.59%) endodermal cells at stage 23 (Figure 1D). Despite absolute numeric differences, both species display comparable relative cellular distributions and volumetric proportions across development (Figure 1E, F). At stage 11.5, the ectoderm comprises the largest cell fraction in both *X. laevis* (64.52% ± 3.14%) and *X. tropicalis* (52.30% ± 1.92%; Figure 1E), whereas volumetric contributions show significant species-specific variation, with ectoderm dominant in *X. laevis* (48.80% ± 7.30%) and endoderm accounting for 34.53% ± 3.26% of embryo volume in *X. tropicalis* (Figure 1F). By stage 23, tissue proportions fully align across species with no statistically significant differences (p > 0.05), where the endoderm constitutes the largest volumetric fraction in both *X. laevis* (41.31% ± 4.67%) and *X. tropicalis* (33.10% ± 3.23%; Figure 1F).

Collectively, these data demonstrate that despite marked differences in absolute cell number and embryo size, and germ layer cell proportions dynamics, *X. laevis* and *X. tropicalis* converge on conserved tissue proportions by tailbud stage 23, indicating a robustly preserved developmental scaling across species.

### 2. Global Interspecies Scaling and Cellular Volume Dynamics

To evaluate how *Xenopus laevis* and *Xenopus tropicalis* compare in overall body architecture, we first performed a global quantitative analysis of embryo volume, total cell number, and cell density at the onset of gastrulation (Stage 11.5) and at the tailbud stage (Stage 23) (Figure 2A–H). During gastrulation, *X. laevis* embryos exhibited a significantly larger physical volume (0.406 mm³ vs. 0.204 mm³; 1.99-fold, p = 0.0028) and higher total cell count (54,505 vs. 21,506 cells; 2.53-fold, p < 0.001) compared to *X. tropicalis*, resulting in a 1.28-fold higher cell density (135,493 vs. 105,482 cells/mm³, p = 0.0234) (Figure 2A–D). By the tailbud stage, *X. laevis* maintained a greater overall volume (0.437 mm³ vs. 0.181 mm³; 2.41-fold, p < 0.001) and higher total cell count (72,320 vs. 42,626 cells; 1.70-fold, p < 0.001) (Figure 2E, F, H). However, density dynamics diverged sharply: *X. tropicalis* displayed a 1.43-fold higher cell density relative to *X. laevis* (236,662 ± SD vs. 165,573 ± SD cells/mm³, respectively; n = 3, p = 0.0018) (Figure 2G, H). This discrepancy indicates that while *X. laevis* undergoes significant volumetric expansion, *X. tropicalis* adopts a distinct strategy of body organization by increasing cellular packing density within a smaller organismal volume. To further determine whether these global differences in volume, cell number, and density reflect uniform or tissue-specific developmental shifts, we examined these metrics across individual germ layers and their derived sub-lineages (Figure S3A–F). At Stage 11.5, *X. laevis* exhibited significantly larger germ-layer volumes than *X. tropicalis* across all primary lineages, most prominently in the mesoderm (*p* < 0.0001) ( Figure S3A). This volumetric excess was accompanied by significantly higher cell numbers in both the ectoderm (*p* < 0.01) and endoderm (*p* < 0.01) (Figure S3B). Surprisingly, local cell packing density at gastrulation did not follow a uniform pattern: while *X. laevis* displayed a significantly higher density within the mesoderm (*p* < 0.05), *X. tropicalis* exhibited a higher cell packing density in the endoderm (*p* < 0.01) (Supplementary Figure 3C). By Stage 23, while total germ-layer volumes remained consistently larger in *X. laevis*—with significant differences persisting in the endoderm and somites (*p* < 0.05)—and cell counts were elevated in the endoderm (*p* < 0.01) and epiderm (*p* < 0.05) (Figure S3D, E), local tissue density homogenized across all sub-lineages, showing no statistically significant differences between species in any individual derived tissue (Figure S3F). Together, these spatial profile analyses reveal that species-specific differences in embryo organization are strongly compartmentalized during gastrulation but converge toward comparable tissue-level packing densities across homologous sub-lineages by the tailbud stage.

**Figure 2.**
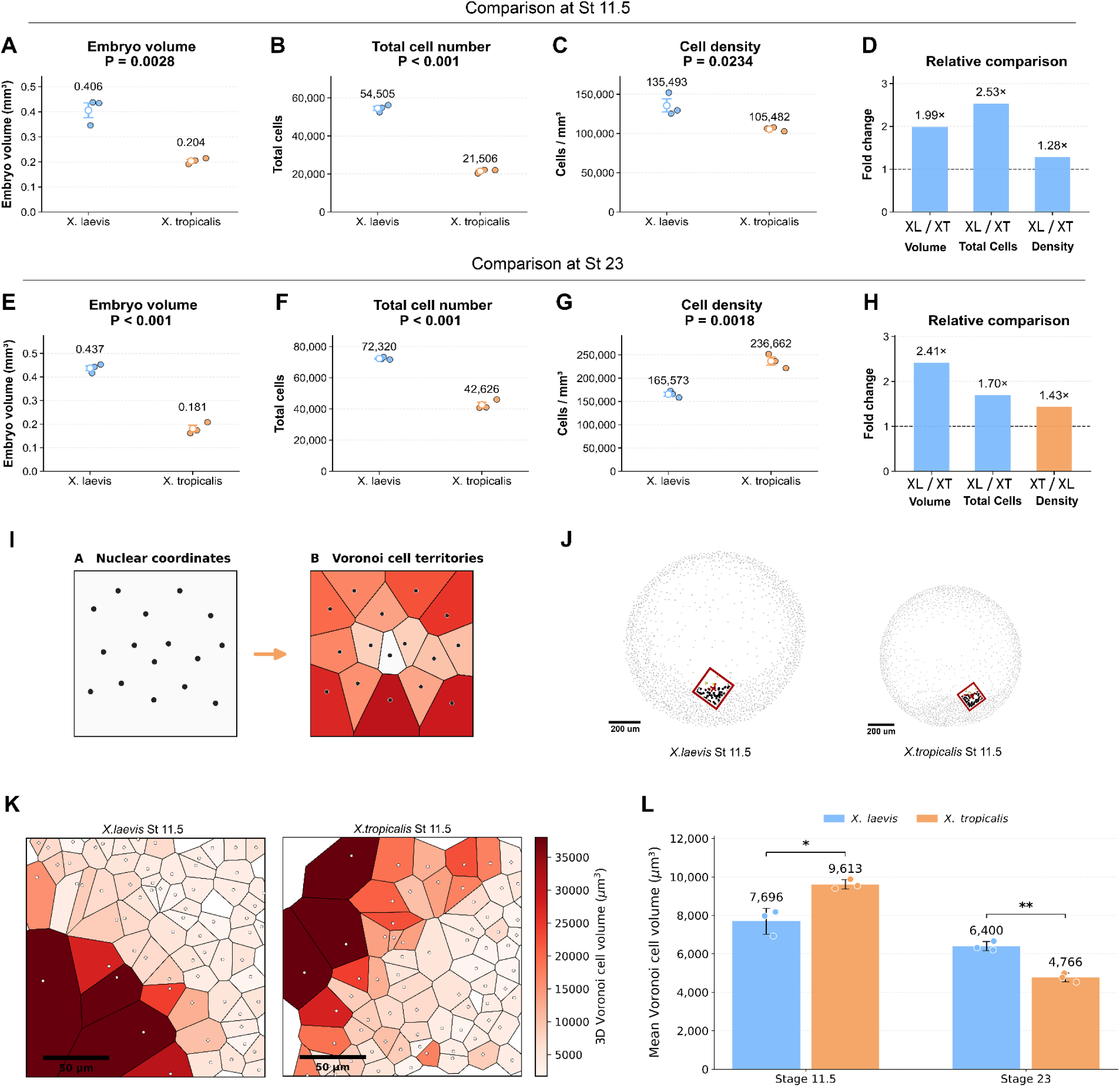
Interspecies volumetric scaling and cellular packing density shifts are driven by dynamic mean cell volume changes in *Xenopus*. **(A–D)** Global architectural comparisons during gastrulation (Stage 11.5) showing **(A)** total embryo volume (mm³), **(B)** total cell count, **(C)** cellular packing density (cells mm³), and **(D)** relative fold-change ratios between *X. laevis* (XL) and *X. tropicalis* (XT). **(E–H)** Global architectural comparisons at the tailbud stage (Stage 23) displaying **(E)** total embryo volume, **(F)** total cell count, **(G)** cellular packing density, and **(H)** relative fold-change ratios, highlighting a density inversion in tailbud *X. tropicalis*. **(I)** Schematic representation of the 3D Voronoi tessellation pipeline, converting discrete nuclear centroids into convex polyhedral spatial territories representing individual cell domains. **(J)** Selection of equivalent anatomical regions in *X. laevis* and *X. tropicalis* gastrulae for micro-scale spatial analysis. **(K)** High-magnification 3D Voronoi tessellations of the selected regions in (J), color-coded by volume (µ*m*^3^), illustrating high cell-size heterogeneity in *X. laevis* compared to a more uniform, medium-sized cell distribution in *X. tropicalis*. **(L)** Quantification of global mean Voronoi cell volumes (µ*m*^3^) across developmental stages, demonstrating an inversion in average cell size between Stage 11.5 and Stage 23. Data represent mean ±SD (n = 3 embryos per species/stage); statistical significance evaluated by unpaired t-test (*p < 0.05,p < 0.01, ***p < 0.001).

To determine how individual cell volumes and physical dimensions scale across development and underpin these density shifts, we implemented 3D Voronoi tessellation based on segmented nuclear coordinates (Figure 2I–K). In this geometric partitioning framework, the 3D space around each nucleus is divided into convex polyhedral territories representing the spatial domain closest to each centroid (Figure 2I), providing a Voronoi-based measure of local 3D cellular density across embryonic regions. To directly compare tissue-level organization between species, two equivalent anatomical regions were selected in *X. laevis* and *X. tropicalis* gastrulae (Figure 2J) and subjected to Voronoi tessellation (Figure 2K). Regional visual inspection in Figure 2K reveals a distinct contrast in cell size heterogeneity at Stage 11.5: *X. laevis* displays a broader mixture of very large and small cells, whereas *X. tropicalis* exhibits a more uniform distribution dominated by medium-sized cells.

This structural divergence in size distribution becomes quantitative when evaluating mean global cell volumes (Figure 2L) and relative cell-size frequency distributions (Figure S3). During gastrulation (Stage 11.5), *X. laevis* cells exhibited a mean Voronoi volume of 7,696 ± 500 µ*m*^3^, whereas *X. tropicalis* cells were larger on average (9,613 ± 200 µ*m*^3^, p < 0.05; Figure 2L), a trend reinforced by the broader tail toward larger individual cell volumes in *X. tropicalis* (Figure S3, Stage 11.5). As development progressed to Stage 23, rapid cleavage and tissue reorganization inverted this relationship: *X. laevis* cells retained a larger average volume (6,400 ± 400 µ*m*^3^) compared to 4,766 ± 300 µ*m*^3^ in *X. tropicalis* (p < 0.01; Figure 2L). The corresponding volume frequency histograms at Stage 23 highlight a sharp leftward shift in *X. tropicalis*, with over 40% of its cells occupying the smallest volume bracket (< 2,500 µ*m*^3^) compared to ∼ 27% in *X. laevis* (Figure S3, Stage 23). Together, these volumetric estimations demonstrate that *X. tropicalis* achieves elevated tailbud cell density through a distinct morphogenetic scaling strategy, accommodating a greater number of smaller cells within a constrained spatial domain.

### 3. Conservation of Spatial Cellular Architecture: Packing, Order, and Geometry

To assess spatial cellular organization during embryonic development, we extracted nuclear coordinates and applied Voronoi 3D spatial tessellations to mathematically define local spatial domains and nuclear neighborhoods as proxies for cell volume and packing (Figure 3A). Using this consolidated approach, we characterized 14 individual morphometric parameters of cellular organization, including volumetric regularity, cell contact, isotropy, anisotropy, local volume, and neighbor variance (Figure S4–S8 for *X. laevis*; Figure S9–S12 for *X. tropicalis*). To enable macro-scale structural tracking, these 14 continuous parameters were equally weighted and integrated into four functional composite classes: spatial order, which quantifies local variance via volumetric regularity, volume variance, neighbor distance variance, and neighbor count variance; cell packing, which measures local crowding through density, mean neighbor number, cell contact area, and neighbor distance; cell geometry, which captures domain scale through local volume, geometry, and contact area; and directionality, which evaluates spatial tensor alignment through orientation, vector alignment, anisotropy, and isotropy (Figure 3B; detailed tissue-level heatmaps provided in (Figure S8 and S12). To facilitate open-access exploration of our 3D dataset, we developed a WebGL-based online visualization platform. The platform includes an interactive viewer (https://xenopus.hms.harvard.edu/3d_models.html) displaying all 14 cellular packing and geometry metrics.

**Figure 3.**
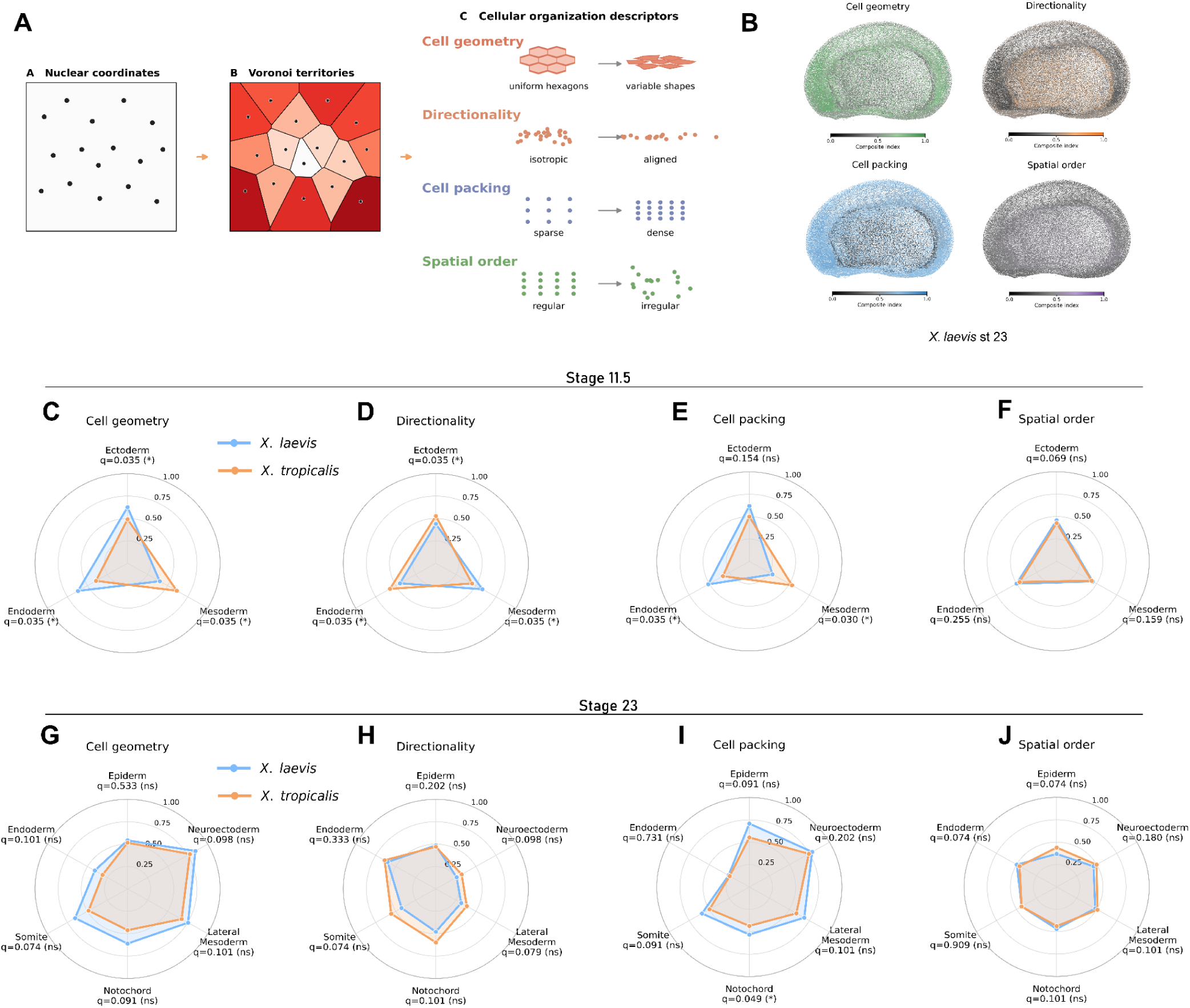
Multi-parametric cellular architecture profiles reveal early germ-layer divergence followed by tissue-level convergence at the tailbud stage. **(A)** Methodological workflow illustrating the extraction of 3D Voronoi cellular territories from segmented nuclear coordinates to define 14 individual organizational descriptors grouped into four functional composite classes: Cell geometry, Directionality, Cell packing, and Spatial order. **(B)** Representative full-embryo 3D point-cloud models (*Xenopus laevis*, Stage 23) color-coded by normalized composite indices for Spatial order, Cell geometry, Cell packing, and Directionality. Full interactive 3D datasets for all 14 quantitative morphometric descriptors and 4 composite architectural indices across all developmental stages and species can be rendered and explored in the online model gallery at https://xenopus.hms.harvard.edu/3d_models.html. **(C–F)** Radar plots comparing tissue-level architectural indices during gastrulation (Stage 11.5) between *X. laevis* (blue) and *X. tropicalis* (orange) across primary germ layers for **(C)** Spatial order, **(D)** Cell packing, **(E)** Cell geometry, and **(F)** Directionality, showing significant early divergence in mesodermal and ectodermal parameters (q < 0.035). **(G–J)** Radar plots comparing tissue-level architectural indices at the tailbud stage (Stage 23) across detailed derived tissues for **(G)** Spatial order, **(H)** Cell packing, **(I)** Cell geometry, and **(J)** Directionality, demonstrating structural convergence between species with no significant differences across most tissue lineages (q > 0.05). Data represent tissue mean values (n = 3 embryos per species/stage); FDR-adjusted statistical significance evaluated by multiple t-tests with Benjamini-Hochberg correction (ns = not significant, *q < 0.05).

Analysis of these architectural patterns across *Xenopus laevis* morphogenesis revealed that during gastrulation (St. 11.5), the embryo exhibits high overall structural uniformity across its primary germ layers without the strong predominance of any single composite class (Figure S8A). As development proceeds through neurulation (St. 15), distinct regional specializations emerge: the neuroectoderm displays markedly elevated levels of cell packing and cell geometry, particularly within its anterior domain, in sharp contrast to the endoderm, which exhibits the lowest values for these same classes (Figure S8B). By the early tailbud stage (St. 23), these tissue-specific cellular architecture signatures become further accentuated, reinforcing the high packing and geometric complexity of the neuroectoderm relative to the loosely packed endoderm (Figure S8C). Multi-parametric stage comparisons across *X. laevis* development reveals no statistically significant differences across stages for any of the composite classes (p > 0.05; Figure S8D), indicating that spatial order, cell packing, cell geometry, and directionality remain overall stable during these developmental transitions.

Having established the spatiotemporal profile in *X. laevis*, we next analyzed the architectural dynamics of *Xenopus tropicalis* (Figure S12). During gastrulation (St. 11.5), intermediate organizational levels were observed across most germ layers (Figure S12A). However, in contrast to *X. laevis*, *X. tropicalis* gastrulae exhibited higher cell packing and cell geometry indices within the mesoderm (Figure 3C-F and Figure S12A), revealing an early structural divergence between the two species during early gastrulation. By the tailbud stage (St. 23), *X. tropicalis* tissues retained a profile closely matching *X. laevis*, with the neuroectoderm exhibiting the highest cell packing and cell geometry indices and the endoderm displaying the lowest (Figure 3G-J and Figure S12B). Temporal comparison between stages in *X. tropicalis* indicated that the largest variations occur in cell geometry and directionality (p < 0.05; Figure S12C)—a trend opposite to that of *X. laevis*: tailbud embryos (St. 23) displayed significantly higher cell geometry scores, whereas gastrulae (St. 11.5) exhibited higher mean directionality (Figure S12C).

Finally, direct interspecies comparative analysis between *X. laevis* and *X. tropicalis* across germ layers demonstrated that during gastrulation (St. 11.5), embryos display significant differences across several tissue-resolved architectural indices, notably in mesodermal and ectodermal cell geometry and directionality (q < 0.035; Figure 3C–F). By the tailbud stage (St. 23), however, most tissue-specific architectural differences were no longer detected across spatial order, packing, geometry, and directionality (q > 0.05; Figure 3G–J), indicating a broad structural convergence between species following morphogenesis.

Together, these quantitative comparisons reveal an intriguing evolutionary pattern: during gastrulation, *X. laevis* and *X. tropicalis* differ substantially not only in absolute cell number and tissue volume, but also in their tissue-level cellular organization. By contrast, by the tailbud stage, fractional tissue proportions and relative layer volumes align between species (Figure 1E, F), with homologous tissues converging toward comparable cellular architectures despite persistent differences in total embryo size and cell density.

### 4. Dividing Cells: Spatial Distribution and Constancy Across Stages and Species

To understand the cellular dynamics governing embryonic scaling, we first evaluated the overall expansion of absolute cell numbers from gastrula (St. 11.5) to early tailbud (St. 23) stages across both species. Across this developmental window, *X. laevis* expands its total cell population from 54,505 ± 1,500 to 72,320 ± 2,100 cells, a net increase of 32.7% (Figure 2B, F). In contrast, *X. tropicalis* undergoes a near-doubling of its cell number, increasing by 98.2% from 21,506 ± 1,000 to 42,626 ± 1,500 cells (Figure 2B, F). This marked difference highlights the smaller *X. tropicalis* embryo engages in a substantially higher relative proliferative expansion during morphogenesis compared to *X. laevis*, driving its pronounced increase in cellular density within a constrained physical volume (Figure 2G, H).

To determine the spatial distribution and physical microenvironments where these active divisions occur, we developed a dedicated machine-learning classification model capable of identifying sister nuclei in an anaphase-like configuration using 3D nuclear spatial coordinates and morphological descriptors (Figure 4A and Supplementary Table 1). Candidate nuclear pairs were evaluated across 10 feature domains, including pair proximity, size similarity, shape/axis orientation, mirror symmetry, interface intensity, and three-level geometry, to robustly distinguish true sister nuclei undergoing active separation from non-dividing adjacent cells (Figure 4A and Supplementary Table 1). Validation against expert-annotated 3D datasets confirmed exceptional classification performance, achieving an overall F1-score of 97.0% at Stage 23 (100% precision, 94.0% recall), as well as 100% F1-score at Stage 11.5 and 88.0% at Stage 15 (Figure S13A–C).

**Figure 4.**
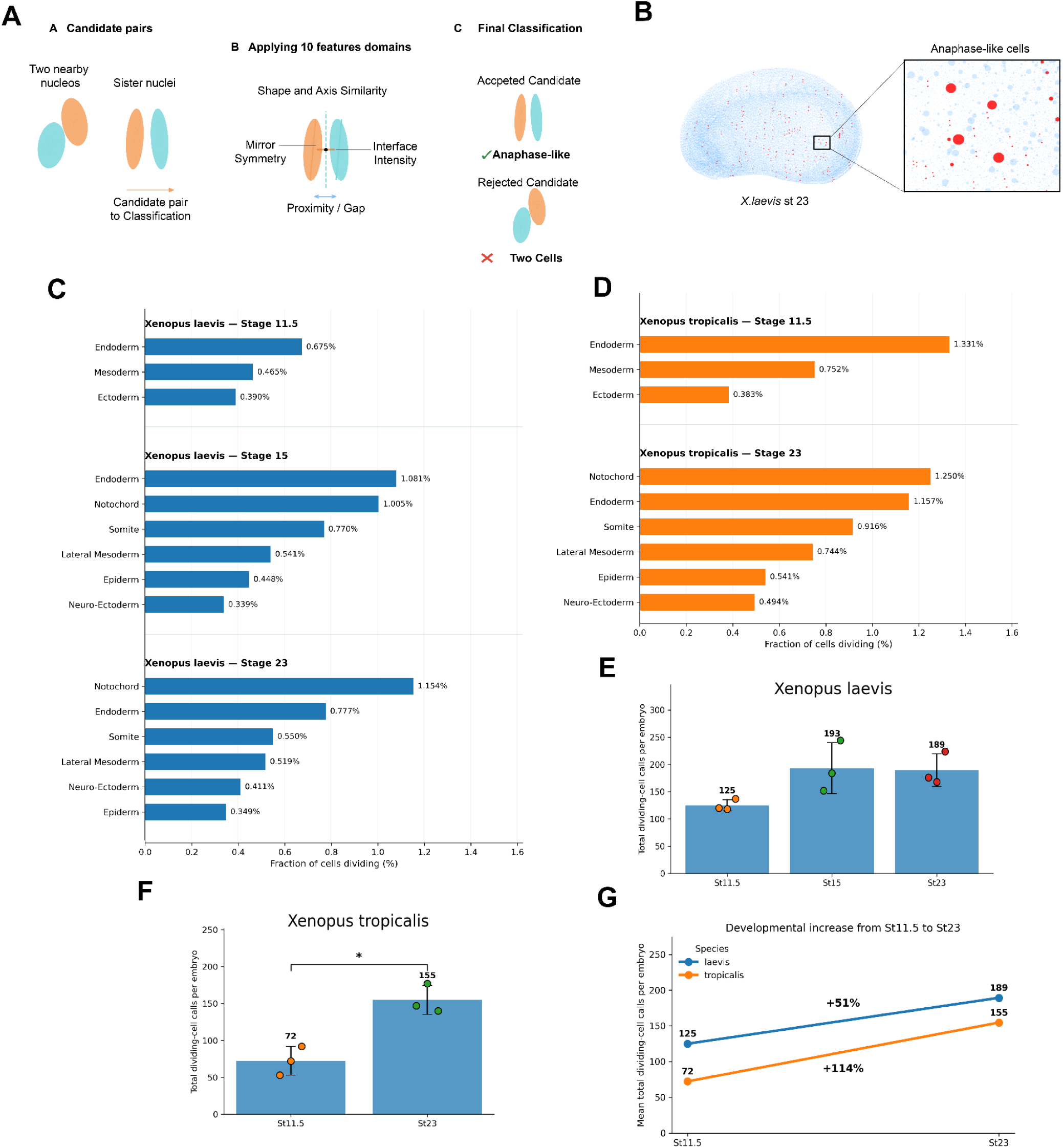
Machine-learning detection of anaphase-like events reveals lineage-specific mitotic dynamics and accelerated proliferative expansion in *Xenopus tropicalis*. **(A)** Schematic of the machine-learning classification pipeline used to evaluate candidate nuclear pairs across 10 morphological and spatial feature domains (e.g., proximity, shape/axis similarity, mirror symmetry, interface intensity) to accept true sister nuclei in anaphase-like configurations and reject adjacent non-dividing pairs. **(B)** Representative full-embryo 3D point-cloud reconstruction (*Xenopus laevis*, Stage 23) showing spatial mapping of detected anaphase-like events (red points) inset within the total nuclear population. **(C–D)** Mitotic index analysis displaying the percentage fraction of cells undergoing division relative to total cell count across primary germ layers and derived lineages for (C) *X. laevis* (Stage 11.5, Stage 15, and Stage 23) and (D) *X. tropicalis* (Stage 11.5 and Stage 23). **(E–F)** Absolute numbers of detected anaphase-like events per embryo across developmental stages in (E) *X. laevis* (125 ± 14 at St 11.5; 193 ± 41 at St 15; 189 ± 23 at St 23; Quasi-Poisson regression p > 0.10) and (F) *X. tropicalis* (72 ± 20 at St 11.5 vs. 155 ± 28 at St 23; *p = 0.033). **(G)** Interspecies comparison of mean total dividing-cell calls from Stage 11.5 to Stage 23, illustrating a +114% proliferative surge in *X. tropicalis* compared to a +51% increase in *X. laevis*. Data represent mean ± SD (n = 3 embryos per species/stage); statistical significance evaluated by unpaired t-test or Quasi-Poisson regression (*p < 0.05).

Applying this validated pipeline allowed full-embryo 3D spatial mapping of active mitotic events across germ layers in both species (Figure 4B and Figure S14). In *Xenopus laevis*, detected anaphase-like pairs remained relatively steady across development (125 ± 14 cells at St. 11.5; 193 ± 41 at St. 15; 189 ± 23 at St. 23; Quasi-Poisson regression p > 0.10; Figure 4E, G). In contrast, *Xenopus tropicalis* exhibited a statistically significant surge in active cell divisions, rising from 72 ± 20 cells at St. 11.5 to 155 ± 28 cells at St. 23 (p = 0.033; Figure 4F, G), representing a 114% increase compared to a modest 51% expansion in *X. laevis* (Figure 4G). Across lineages, both species displayed comparable proportional trends, with high mitotic fractions within the ectoderm at early stages and endoderm/notochord in tailbud stages (Figure 4C, D and Figure S14).

Despite differences in absolute cell counts, calculating the mitotic index, the fraction of dividing cells relative to the total nuclear population, revealed that both species maintain a conserved proportional allocation of dividing cells across physical scales (∼0.2%–0.4% instantaneous anaphase-like fraction; Figure 4C, D). *X. tropicalis*, however, maintained a persistently higher fractional mitotic activity (∼1.4-fold higher than *X. laevis* at equivalent stages), directly fueling its pronounced cell density surge within a smaller physical volume (Figure 2G, H and Figure 4G). This conserved scaling aligns with classical observations in *Xenopus* (Junidi et al., 1983; Cooke, 1979a) and supports findings in hybrid embryos where intrinsic cell-cycle control governs developmental timing independently of body scale (Gibeaux et al., 2018). Furthermore, the elevated division rate in *X. tropicalis* is consistent with its faster overall developmental rate compared to *X. laevis* (Khokha et al., 2002), with both species exhibiting progressive lengthening of the cell cycle during tissue differentiation (Thuret et al., 2015).

Finally, we evaluated how these mapped mitotic events correlate with local microenvironmental cellular architecture (Figure 5). Spearman rank correlation analysis between anaphase-like occurrence and composite organizational classes revealed that cell divisions occur non-uniformly, concentrating in spatial domains characterized by high spatial order and high directionality (Figure 5A).

**Figure 5.**
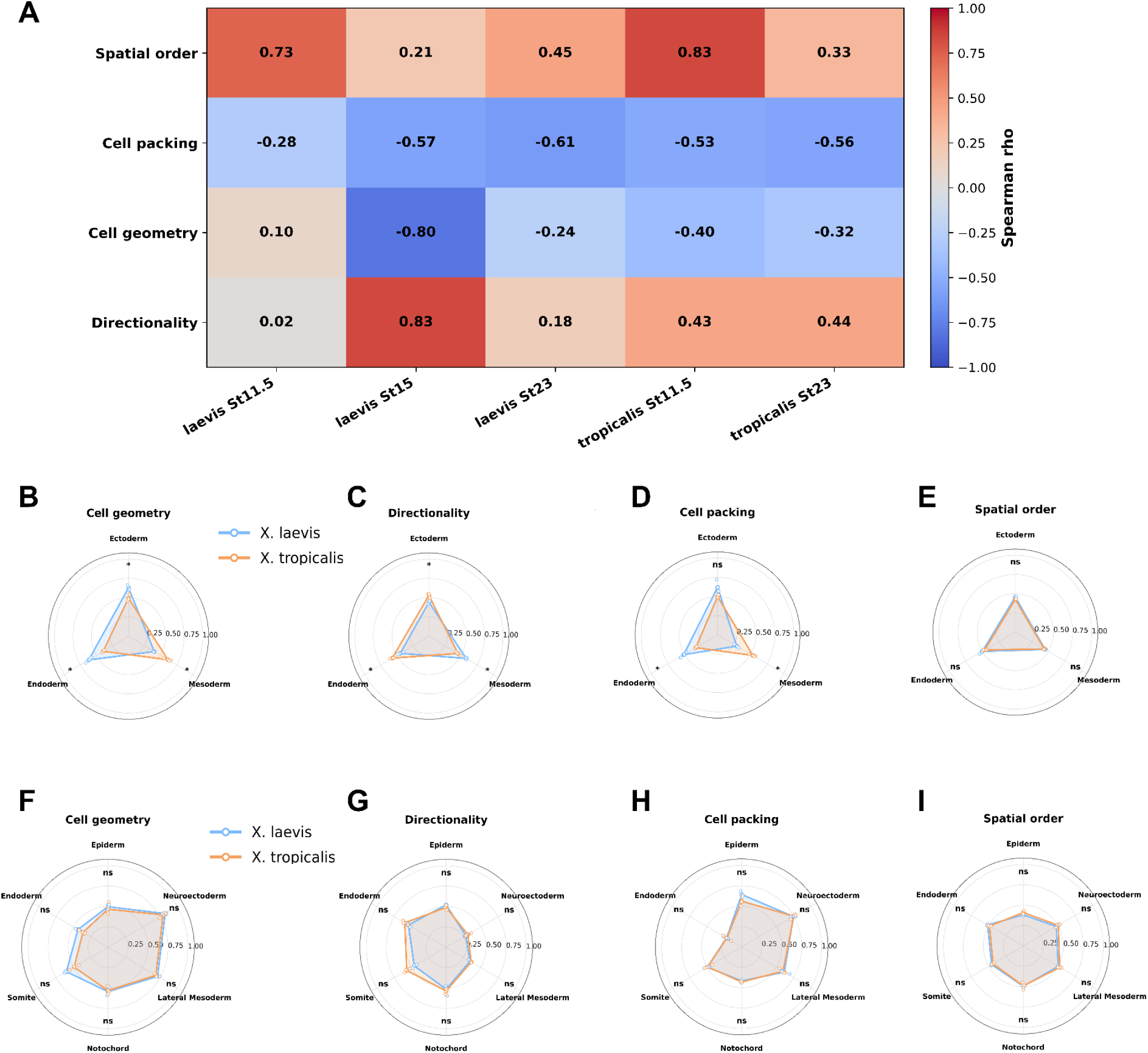
Spatial coupling of active cell divisions to microenvironmental cellular architecture demonstrates progressive interspecies convergence during morphogenesis. **(A)** Heatmap displaying Spearman rank correlation coefficients (⍴) between active anaphase-like occurrences and four composite architectural classes (Spatial order, Cell packing, Cell geometry, and Directionality) across developmental stages in *Xenopus laevis* (St 11.5, St 15, St 23) and *Xenopus tropicalis* (St 11.5, St 23), showing strong positive coupling to Spatial order and Directionality, and negative correlation with Cell packing. **(B–E)** Radar plots comparing microenvironmental architectural properties of mitotic regions during gastrulation (Stage 11.5) between *X. laevis* (blue) and *X. tropicalis* (orange) across primary germ layers for **(B)** Spatial order, **(C)** Cell packing, **(D)** Cell geometry, and **(E)** Directionality. **(F–I)** Radar plots comparing microenvironmental architectural properties of mitotic regions at the tailbud stage (Stage 23) across detailed derived tissues for **(F)** Spatial order, **(G)** Cell packing, **(H)** Cell geometry, and **(I)** Directionality, highlighting architectural convergence in proliferative microenvironments across all tissue lineages (q > 0.05). Data represent mean composite indices (n = 3 embryos per species/stage); FDR-adjusted statistical significance evaluated by multiple t-tests with Benjamini-Hochberg correction (ns = not significant).

When comparing gastrula embryos (St. 11.5), the mesoderm emerged as the most divergent tissue between *X. laevis* and *X. tropicalis* regarding proliferative microenvironments, whereas ectoderm and endoderm differed only modestly in cell packing and spatial order (Figure 5B–E). By contrast, at the tailbud stage (St. 23), both species converged on the same structural distribution of cell divisions across all tissue lineages, showing no statistically significant architectural differences in proliferative zones (q > 0.05; Figure 5F–I).

In summary, while *X. laevis* and *X. tropicalis* exhibit species-specific tissue architecture and proliferative microenvironments during gastrulation (Figure 3C–F and Figure 5B–E), morphogenesis drives a progressive structural convergence, yielding equivalent architectural and mitotic organization by the tailbud stage (Figure 3G–J and Figure 5F–I).

## DISCUSSION

### Developmentally dynamic scaling of cell number, density, and cellular space

Our whole-embryo quantitative analysis reveals that scaling between *Xenopus laevis* and *Xenopus tropicalis* cannot be explained by a fixed proportional relationship among embryo size, cell number, and cellular density. Instead, these parameters are developmentally dynamic, and their relationships change substantially between gastrulation and the early tailbud stage. Although *X. laevis* is consistently larger and contains more cells than *X. tropicalis*, the manner in which these cells occupy embryonic space changes over developmental time.

This distinction is particularly evident when cell density is considered. At St. 11.5, *X. laevis* displays a higher cellular density than *X. tropicalis*, whereas by St. 23 this relationship is reversed, with *X. tropicalis* reaching approximately 1.43-fold higher density despite remaining substantially smaller in total volume and lower in absolute cell number. Thus, high cell density is not an intrinsic or developmentally invariant characteristic of *X. tropicalis*. Rather, the two species follow different trajectories of cellular packing that cross during morphogenesis.

The Voronoi-based analysis reinforces this interpretation. At gastrulation, the mean spatial domain associated with each nucleus is larger in *X. tropicalis* than in *X. laevis* (9,613 versus 7,696 µm³), consistent with the lower cellular density of *X. tropicalis* at this stage. By St. 23, this relationship is inverted: *X. tropicalis* exhibits substantially smaller Voronoi domains than *X. laevis* (4,766 versus 6,400 µm³), accompanying its higher cellular density. These reciprocal changes indicate that interspecies scaling is not simply established by initial differences in egg or cell size but is progressively remodeled during embryogenesis (Miller et al., 2023).

Differences in egg size, genome content, and nuclear scaling are likely to contribute to the initial physical conditions of the two species (Levy & Heald, 2010; Gibeaux et al., 2018). However, the developmental inversion observed here indicates that these initial conditions alone cannot explain the cellular organization present at later stages. Morphogenesis progressively modifies the relationship among cell number, tissue volume, and the amount of physical space available per cell. In this sense, embryonic scaling appears to be an active developmental process rather than the passive preservation of a size relationship established at fertilization.

This interpretation is consistent with classical observations showing that vertebrate embryos can preserve tissue organization despite major experimentally induced changes in cell size or cell number (Fankhauser, 1945; Fankhauser, 1972). Our results extend this principle to a natural interspecies comparison and suggest that robustness of embryonic form can coexist with substantial flexibility in the cellular parameters used to construct that form.

### Transient divergence and later convergence of tissue allocation

The comparison of germ-layer composition further demonstrates that scaling is not achieved through invariant proportional allocation throughout development. At St. 11.5, *X. laevis* and *X. tropicalis* differ significantly in the relative allocation of cells to the ectoderm and mesoderm, whereas the endoderm is comparatively similar. Therefore, the earliest stages analyzed here already show that equivalent developmental stages do not necessarily contain identical proportional cellular compositions.

By St. 23, however, this situation changes markedly. Despite substantial differences in total embryo volume, cell number, and cellular density, no significant interspecies differences are detected in the relative cellular representation of the major differentiated tissue compartments analyzed. A similar pattern is observed when the proportional volume occupied by these tissues is compared. The data therefore support a model of developmental convergence rather than continuous proportional invariance.

This distinction has important implications for understanding developmental scaling. Rather than requiring identical cell allocation from the beginning of morphogenesis, the developmental programs of the two species appear capable of accommodating early quantitative differences while subsequently reaching broadly comparable tissue proportions. The conserved outcome therefore does not require a conserved trajectory.

Such behavior is conceptually consistent with studies showing that embryonic patterning systems can adjust tissue proportions after perturbations in embryo size or cell number (Almuedo-Castillo et al., 2018). Our data do not directly identify the molecular mechanisms responsible for the convergence observed here. Instead, they show that the integrated developmental system is sufficiently flexible to generate similar later tissue allocation from quantitatively different earlier cellular states.

The increase in mesodermal representation observed during development is also compatible with the extensive morphogenetic rearrangements and recruitment processes previously described in *Xenopus* gastrulation and neurulation (Cooke, 1979a, 1979b). Because the present study compares fixed embryos at discrete developmental stages rather than performing lineage tracing, however, these changes cannot be attributed directly to specific cell recruitment trajectories. They should instead be interpreted as population-level changes consistent with established developmental processes.

### A conserved organizational framework with flexible cellular implementation

The morphometric analysis provides a quantitative basis for distinguishing conservation from species-specific cellular strategies during development. By integrating local neighborhood measurements into composite indices of spatial order, cell packing, cell geometry, and directionality, we find that *X. laevis* and *X. tropicalis* exhibit temporal dynamics characterized by early architectural divergence followed by late structural convergence.

During gastrulation (St. 11.5), the two species display distinct cellular architectures and tissue-specific organizational profiles. Significant differences emerge across global morphometric indices, cell geometry, and regional packing, particularly within the mesoderm and ectoderm, despite early overall similarities in macro-scale spatial symmetry. This early divergence occurs alongside marked disparities in absolute cell number, total embryonic volume, cellular density, and germ-layer allocation. Thus, the initial cellular paths and local tissue organizations employed by *X. laevis* and *X. tropicalis* to undergo gastrulation are inherently species-specific.

The situation at St. 23 reveals a striking developmental transition. By this early tailbud stage, homologous tissues across both species broadly converge in their relative cellular composition, volume proportions, and local morphometric profiles. Despite persistent differences in absolute embryo scale and an inversion in cellular density, tissue-level packing, orientation, and spatial geometry become broadly equivalent between *X. laevis* and *X. tropicalis*. Consequently, macro-scale tissue architecture and local cellular parameters undergo progressive alignment during morphogenesis.

This temporal dynamic demonstrates that developmental conservation in *Xenopus* operates through convergent trajectories rather than persistent cellular invariance. Early embryonic stages permit significant flexibility and species-specific variation in cell allocation, geometry, and local packing. As morphogenesis proceeds, these distinct initial configurations are actively remodeled to converge upon a highly conserved, equivalent tissue architecture at tailbud stages. The conserved property of vertebrate embryogenesis may therefore reside less in maintaining identical cellular parameters from the outset than in guiding divergent early cellular states toward a shared higher-order organizational archetype.

This interpretation refines the concept of an embryonic “cellular blueprint.” The blueprint should not be understood as a rigid template requiring identical initial cell counts, densities, or micro-architectures across species. Instead, our data support a constrained developmental framework that accommodates species-specific cellular implementations during early cleavage and gastrulation while ensuring convergence toward invariant tissue-level organization. Such a model aligns with the emerging concept of organizational archetypes and conserved morphogenetic frameworks, in which variable cell-level distributions across early development collapse into shared higher-order spatial organization states as body plan specification completes (Morishita et al., 2023; Brambach et al., 2025).

This cellular-level pattern parallels an independent finding at the transcriptomic level in same two species: cross-species bulk gene expression divergence is concentrated in the earliest developmental stages and declines steadily thereafter, a trend attributed to progressive stabilization of cell-fate-determination transcriptional programs consistent with a developmental hourglass model (Yanai et al., 2011). The convergence in cellular architecture we observe by St. 23 may represent the morphological correlation of this same underlying canalization (Waddington, 1942) of gene regulatory programs governing cell identity. Notably, Yanai et al. (2011) raised the possibility that transcriptome-level convergence could partly reflect increasing cell-type diversity diluting bulk RNA measurements over developmental time, rather than genuine tissue-level convergence. Because our architectural indices are computed at single-tissue resolution rather than as whole-embryo averages, they are not subject to this confound, and thus provide direct, cell-resolved evidence that the convergence in question is a real property of homologous tissues rather than an artifact of increasing sample heterogeneity.

The physical mechanisms underlying this architectural convergence remain to be directly elucidated. While the progressive alignment of local architecture likely involves shared tissue mechanics, differential cell adhesion, and cortical tension constraints (Heisenberg & Bellaïche, 2013), our morphometric parameters capture spatial relationships derived from nuclear positions. They identify a clear developmental transition from early cellular divergence to late architectural equivalence, providing a quantitative framework for future mechanistic and biophysical exploration.

### Distinct proliferative trajectories contribute to developmental remodeling

The spatial analysis of anaphase-like nuclear pairs adds another dimension to this developmental comparison. In *X. laevis*, the number of detected dividing pairs does not change significantly among St. 11.5, St. 15, and St. 23 when total nuclear number is taken into account. In *X. tropicalis*, in contrast, anaphase-like events increase significantly between St. 11.5 and St. 23. Descriptively, this corresponds to an approximately 114% increase in *X. tropicalis*, compared with an approximately 51% increase in *X. laevis*.

These trajectories are consistent with greater developmental remodeling of cellular packing in *X. tropicalis* between gastrulation and St. 23. However, the interspecies interaction testing whether the developmental increase itself differs between species did not reach statistical significance. Thus, the data support a significant developmental increase within *X. tropicalis* that should be interpreted as a qualitative trend rather than as evidence for a formally distinct species-specific proliferation program.

Similarly, the association between increased anaphase-like activity and the high cellular density of St. 23 *X. tropicalis* should not be interpreted as demonstrating direct causation. Cellular density is determined jointly by cell number and the physical volume occupied by the tissue, and the Voronoi analysis shows that the average spatial domain surrounding individual nuclei also decreases strongly during *X. tropicalis* development. The observed increase in anaphase-like events, decrease in local cellular space, and increase in density are therefore mutually consistent components of the same developmental transition, but the present analysis cannot establish which of these changes is causally upstream of the others.

Importantly, the spatial environment associated with cell division is also structured. Across developmental conditions, anaphase-like events preferentially occur in regions characterized by high spatial order and higher directionality, indicating that mitotic events are not randomly distributed throughout embryonic tissue. Spatial mapping demonstrated higher mitotic activity in ventral and lateral domains, whereas dorsal axial structures (notochord and somites) showed reduced proliferation as cells exit the cell cycle to undergo axial elongation and differentiation (Cooke, 1979b). This localized mitotic inhibition is consistent with cell-cycle arrest required during convergent extension in paraxial mesoderm (Leise & Mueller, 2004). Conversely, active mitosis within the gastrulating ectoderm plays an essential inductive role, as oriented cell divisions instruct neuroectoderm anterior-posterior patterning and neural plate regionalization (Velloso et al., 2026). This observation agrees with classical descriptions of spatially heterogeneous proliferative activity during *Xenopus* development (Cooke, 1979a, 1979b).

Recent work on nuclear and spindle positioning in early embryos demonstrates that cell geometry and polarity cues compete to position the mitotic apparatus, with polarity progressively outcompeting geometry as development proceeds (Nommick et al., 2026). Our observation that anaphase-like divisions preferentially occur in regions of high directionality and high spatial order (Figure 5A) is consistent with a model in which local tissue architecture influences, and is influenced by, the biophysical constraints on individual division events.

The interspecies comparison again reveals temporal decoupling. During gastrulation, the mesoderm shows the biggest differences between species in the structural environment associated with cell division, consistent with the broader divergence in tissue architecture at this stage. By St. 23, both the structural properties of proliferative regions and the architecture of homologous tissues become substantially more similar between species. Thus, the organization of proliferative niches and global tissue architecture represent related but distinguishable levels of embryonic organization.

Together, these findings suggest that robust morphogenesis does not require identical spatial or temporal distributions of cell division. Instead, proliferation appears to represent one of several flexible cellular processes through which embryos of different physical scales can reach comparable higher-order tissue organization.

### Conservation of developmental outcome does not imply conservation of developmental trajectory

Taken together, our results support a model in which embryonic scaling emerges from the interaction between constrained developmental outcomes and flexible cellular trajectories. Several observations converge on this interpretation. First, cellular density and Voronoi-derived cellular space undergo opposite interspecies relationships at St. 11.5 and St. 23. Second, germ-layer allocation differs during gastrulation but becomes broadly similar by the early tailbud stage. Third, species-specific differences in cellular architecture and tissue geometry are most pronounced early during gastrulation, whereas local structural parameters progressively align to reach macro-scale equivalence at later stages. Finally, the spatial organization of cell division also transitions from an early species-specific state to a shared structural distribution following morphogenesis

Therefore, conservation and divergence should not be viewed as mutually exclusive explanations of Xenopus scaling. Rather, they occur simultaneously at different biological levels. What appears conserved is the ability of the developmental system to generate a recognizable and proportionally organized body plan. What remains flexible is the combination of cell number, cellular density, local geometry, tissue allocation, and proliferative activity through which this state is achieved.

This distinction may help explain the remarkable robustness of vertebrate development. A developmental system in which every cellular parameter had to scale identically would be highly sensitive to changes in embryo size, genome content, or initial cell dimensions. In contrast, a system in which several cellular variables can compensate for one another while higher-order organizational constraints are maintained would be inherently more robust. The comparison between *X. laevis* and *X. tropicalis* provides a natural example of such flexibility.Identifying the species-specific molecular differences at work should deepen our understanding of signaling mechanisms employed to fine-tune normal tissue development and how their deviations could translate into pathological situations.

### Limitations and future directions

Several limitations should be considered when interpreting these results. First, the Voronoi domains used here represent geometric partitions derived from nuclear centroid positions and should therefore be regarded as proxies for the physical space associated with individual cells rather than direct measurements of membrane-defined cell volume. Likewise, distance-derived measures of local contact and packing quantify spatial relationships among nuclei but do not directly measure cell-cell junctions, adhesion forces, cortical tension, or membrane spatial organization.

Second, the study is based on fixed embryos sampled at discrete developmental stages. It therefore provides high-resolution spatial snapshots but does not track individual cells through time. Changes in tissue composition between stages cannot consequently be assigned directly to specific lineage transitions, cell rearrangements, migration events, or individual proliferative histories.

Third, the cell division classifier specifically identifies sister nuclei displaying an anaphase-like configuration based on spatial symmetry and paired morphology. The number of detected events therefore represents a spatial snapshot of a highly restricted sub-stage of M-phase rather than the total fraction of proliferating cells across the entire cell cycle. While classical mitotic markers such as Phospho-Histone H3 (pH3) label cells throughout the broader duration of mitosis, our spatial organization detection focuses strictly on spatially resolved division events. Consequently, these counts should be interpreted as a relative spatial metric of active physical separation rather than a cumulative measurement of cell-cycle speed or total proliferative output.

Finally, the relatively small number of biological replicates limits the power to distinguish true equivalence from failure to detect a difference. Conditions showing no significant interspecies difference should therefore be interpreted as exhibiting similar values within the resolution of the present dataset rather than as demonstrating strict mathematical invariance. Future analyses combining larger embryo cohorts, live imaging, membrane-level cellular segmentation, lineage tracing, and spatially resolved molecular measurements will be important for determining how the architectural states identified here are generated mechanistically.

## Conclusions

Our whole-embryo 3D analysis demonstrates that developmental scaling between *X. laevis* and *X. tropicalis* is achieved neither by simple proportional multiplication of cell number nor by rigid conservation of every cellular parameter. Instead, embryo size, cell number, cellular density, local spatial domains, tissue allocation, architecture, and proliferative organization follow partially independent developmental trajectories.

Despite these differences, the two species repeatedly converge at higher levels of organization. This suggests that the robustness of the vertebrate body plan may arise from developmental constraints that act on collective tissue organization while allowing substantial flexibility in the cellular strategies used to satisfy those constraints.

Rather than representing a fixed cellular blueprint, embryonic scaling in *Xenopus* therefore appears to rely on a flexible cellular implementation of a constrained developmental architecture. Such flexibility provides a mechanism through which embryos differing substantially in physical scale, genome content, cell number, and cellular organization can nevertheless generate comparable body plans.

## MATERIALS AND METHODS

### Embryo Preparation and Staging

All animal experiments were conducted following the rules and regulations of the Harvard Medical School IACUC (Institutional Animal Care and Use Committee #1365-6). Females of Xenopus laevis were primed and boosted with 10 U and 700 U of HCG (human chorionic gonadotrophin, Chorulon, Merck, Germany), respectively. Testes were collected after male euthanasia. Eggs were laid and collected in 1× MMR, excess medium was removed, and eggs were fertilized in vitro using sperm solution in 1× MMR. Fertilized eggs were dejellied using 2% (w/v) cysteine at pH 7.8 in 0.1× MMR. Embryos were cultured in 0.1× MMR at 18–22 °C and staged according to the Nieuwkoop and Faber (1994) table. X. tropicalis embryos were obtained by in vitro fertilization according to Lane et al. (2022). Embryos from both species at stages 11.5 and 23, as well as X. laevis embryos at stage 15, were included in the quantitative comparative analysis. X. tropicalis stage 15 embryos were processed during the experimental workflow but were not included in the final quantitative dataset. The selected embryos were fixed in MEMFA (0.1 M MOPS, pH 7.4, 2.0 mM EGTA, 1.0 mM MgSO₄) containing 4% (v/v) formaldehyde overnight at 4 °C.

For each species and developmental stage, five embryos were processed in parallel as an initial experimental batch (n = 5). Prior to quantitative image analysis, embryos were assessed exclusively on the basis of predefined technical quality criteria, including preservation of overall morphology, absence of physical damage, and sufficient fluorescence signal-to-noise ratio throughout the imaged volume. Quality assessment and embryo exclusion were performed before examination of the quantitative results and were therefore independent of the biological measurements reported in this study. Embryos that did not meet these predefined imaging-quality criteria were excluded from downstream analysis.

Three embryos per species and developmental stage were retained for quantitative analysis (n = 3). Each biological replicate originated from an independent fertilization, such that the three embryos represented independent biological replicates rather than technical replicates from a single fertilization event. The same predefined quality-control procedure was applied across experimental groups, and no embryo was excluded on the basis of its measured cell number, tissue composition, spatial organization, or other quantitative phenotype. The final dataset therefore comprised three independent biological replicates per species and developmental stage.

### Pigment Removal and Nuclear Staining

Fixed embryos were bleached with PBS containing 5% hydrogen peroxide and 0.1% Tween-20 (v/v) under light to remove pigmentation. To identify the optimal nucleic acid probe for deep-tissue volumetric imaging in opaque Xenopus embryos, we systematically evaluated three fluorescent dyes: SYTOX Green (1:10,000), DAPI (1,µg/mL), and TO-PRO-3 (1µM) in PBS with 0.02% Triton X-100. Comparative optical sectioning and 3D signal penetration analysis revealed that TO-PRO-3 provided superior signal-to-noise ratio and consistent fluorescence detection across deep internal embryo volumes, whereas DAPI and SYTOX Green exhibited reduced penetration depth and weaker signal attenuation in yolk-rich regions. Consequently, TO-PRO-3 was selected as the standard far-red nuclear dye for all whole-embryo quantitative pipelines. Stained embryos were extensively washed with PBS containing 0.02% Triton X-100 (v/v).

### iDISCO Tissue Clearing

After washing, the embryos were dehydrated in H2O/ethanol (v/v; 30%, 50%, 70%, 90%, and 100%) for 24 h. After dehydration, embryos were transferred to ethyl cinnamate (ECi, 100% - Sigma-Aldrich W243019-1KH-K).

### Imaging

Cleared embryos were imaged using a Nikon Ti2 inverted microscope equipped with a Yokogawa CSU-W1 spinning disk confocal system and a Hamamatsu ORCA-Fusion sCMOS camera. Whole-embryo overviews were acquired using a Plan Apo λ 10×/0.45 NA objective, while regional details were captured with a 20×/0.75 NA objective. Volumetric 3D imaging (Z-stacks) was performed with a 2 µm step size (e.g., 480 optical slices for the 10× overview). Fluorescence excitation was achieved using a 640 nm laser, and emission was collected through a Chroma ET705/72m bandpass filter.

### Image Analysis and Segmentation

3D nuclear segmentation was performed using StarDist3D (Weigert et al., 2020). Prior to inference, images were resampled to isotropic voxels (2.0 µm × 2.0 µm × 2.0 µm) and enhanced via a custom percentile-based normalization and gamma correction. To optimize memory, inference was run on overlapping image tiles (15%) using conservative thresholds (probability = 0.30, NMS = 0.20). To validate segmentation accuracy, model predictions were rigorously evaluated against expert-verified ground truth datasets across representative tissue regions (including endoderm, lateral mesoderm, somites, neuroectoderm, and notochord). Objects detected by StarDist3D were individually audited alongside manual annotations to confirm precise boundary detection and instance separation in dense 3D arrangements. Based on this expert-verified benchmark, the automated StarDist3D pipeline demonstrated robust segmentation performance across all developmental stages, achieving an overall F1-score of 98.17% with high precision and recall across embryonic structures.

3D spatial coordinates and volumetric data were subsequently extracted via scikit-image. All image-processing pipelines were implemented in custom Python scripts available at the project repository.

All image-processing and quantitative analyses were performed using custom Python pipelines developed for whole-embryo Xenopus datasets. The workflow includes image tiling, StarDist3D nuclear segmentation, segmentation post-processing, nuclear coordinate extraction, germ-layer classification, Voronoi-based cell-size and density measurements, calculation of local tissue-architecture indices, and machine-learning-based identification of dividing cells. Source code and documentation are publicly available at [https://github.com/mauriciohugox-lab/Xenopus-Segmentation].

### Layer annotation

To identify the embryonic germ layers, manual annotations were performed using the multi-dimensional image viewer Napari. Anatomical and histological references were based on The Early Development of Xenopus laevis: An Atlas of Histology (Hausen & Riebesell, 1991). To optimize the annotation workflow and increase efficiency without compromising segmentation quality, manual annotation was performed on every fourth optical slice, resulting in an 8 µm Z-spacing between annotated planes. This sampling strategy significantly reduced the total number of manually processed frames while maintaining a high-fidelity representation of the tissue architecture. Finally, each annotated layer was exported and saved individually as a separate file for downstream processing.

### Statistical Analysis Method Summary

Statistical comparisons were tailored to the underlying variance structure of the dataset. For direct stage-matched interspecies comparisons within specific tissue lineages (Figures 1–5), sample variances were evaluated for homogeneity, and two-tailed unpaired Student’s t-tests were applied. Conversely, for cross-stage global architectural analyses (Figure S8D), where cellular reorganization introduces variance heteroscedasticity across developmental stages, two-sided Welch’s t-tests (which do not assume equal variance) were utilized alongside Benjamini-Hochberg false discovery rate (FDR-BH) adjustments to correct for multiple testing.

### Cellular architecture mapping

The three-dimensional spatial organization and local geometry of embryonic tissues were quantified from extracted nuclear coordinates (x, y, z). Anisotropic rescaling was first applied to correct for differences between axial and lateral voxel dimensions, placing all coordinates in real physical space (µm). To characterize cellular packing and domain partitioning, a 3D Voronoi tessellation was constructed, from which individual cell volumes (Local Volume) and volumetric variance (Variance of Volume, Volumetric regularity) were derived. In parallel, local spatial neighborhoods were defined within a fixed Euclidean radius, excluding edge-boundary nuclei with insufficient neighbors to prevent boundary artifacts. From these local interactions, density, spatial proximity (Cell contact, calculated as log-transformed inverse squared distances), neighbor distance statistics (Distance from Neighbors, Variance of distance), and coordination metrics (Number of Neighbors, Variance of the number of Neighbors) were computed. Spatial order and directional alignment (Local Orientation, Vector Alignment) were evaluated through radial and angular vector distribution analyses. Additionally, local microenvironment geometry was characterized via Principal Component Analysis (PCA) of the 3D neighbor covariance matrix to extract tissue Isotropy, Anisotropy, and overall Geometry. Finally, to enable rigorous quantitative comparison and heatmap visualization across species and stages, all continuous metrics were standardized using a unified global transformation: metric values were clipped at global 5th and 95th percentiles derived from the pooled dataset across all stages and species, followed by global Min–Max rescaling to a dimensionless 0–1 scale, ensuring identical reference limits were applied to all individual embryos. To synthesize these 14 continuous morphometric measurements into macro-scale structural descriptors, all features were min-max normalized to a dimensionless 0–1 scale following 5th and 95th percentile clipping. To ensure that all variables contributed to each composite in a biologically consistent direction, metrics in which higher values represented greater heterogeneity or reduced packing were inverted as (1-x) after normalization. This transformation was applied to volume variance, neighbor-distance variance, neighbor-count variance, and mean neighbor distance. The features were then grouped with equal weight into four core functional composite classes: (1) Spatial Order, integrating volumetric regularity and the inverted volume, neighbor-distance, and neighbor-count variance metrics; (2) Cell Packing, integrating local density, mean neighbor count, cell-contact proxy, and inverted mean neighbor distance; (3) Cell Geometry, integrating local volume, overall geometry, and cell-contact proxy; and (4) Directionality, integrating local orientation, vector alignment, and anisotropy. Isotropy was retained as an individual morphometric descriptor but excluded from the Directionality composite because it represents the complementary direction of the anisotropy metric. Composite indices were calculated as the unweighted arithmetic mean of their constituent directionally aligned normalized metrics.

### Anaphase-like identification model

To quantitatively map cell division events across embryonic stages, candidate nuclear pairs were generated from 3D centroid coordinates using a density-based spatial clustering algorithm (DBSCAN). Candidate generation was performed in real physical space by scaling the Z-axis according to voxel anisotropy, retaining only clusters of exactly two nuclei to match manual curator selection. Non-mitotic single nuclei (One cell) and ambiguous annotations (Borderline) were excluded, framing the task as a binary classification between dividing cells (Dividing Cell, positive class) and non-dividing adjacent pairs (Two cells, negative class). For each candidate pair, a comprehensive suite of interpretable 3D biological features was extracted across ten distinct functional domains, including pair proximity, volume/size similarity, shape and axis alignment, fluorescence intensity profiles, mirror symmetry, three-level internal axis alignment, interface bridge intensity, composite sister scores, and prototype-derived similarity metrics (Supplementary Table 1). Notably, the three-level geometry evaluated structural coherence by sampling top, middle, and bottom regions along each nucleus’s internal major axis to enforce ordered, non-crossing parallel configurations during anaphase. Model validation was conducted across three distinct developmental stages—Stage 11.5 (Gastrula; 41 positive / 309 negative candidate pairs), Stage 15 (Neurula; 33 positive / 314 negative candidate pairs), and Stage 23 (Tailbud/Larva; 45 positive / 305 negative candidate pairs)—against expert manual annotations (119 positive and 928 negative candidate pairs in total across stages). Classification was performed using a Random Forest classifier (600 trees, max_depth = None, min_samples_leaf = 2, max_features = “sqrt”, class_weight = “balanced_subsample”) trained with leave-one-stage-out validation and calibrated via 3-fold stratified sigmoid Platt scaling. Final candidate pairs were classified as dividing cells using a high-confidence global probability threshold (p≥ 0.93 for primary DBSCAN pairs; p ≥ 0.990 for KNN rescue candidates) combined with strict biological gates (V12B high-confidence filter) requiring spatial proximity, size/intensity matching, and structural symmetry. This framework demonstrated exceptional performance, achieving absolute precision (1.00 across all stages) and robust F1-scores exceeding 0.85.

## Acknowledgments

We acknowledge funding from the Lemann Brazil Research Fund to JGA and LP and from NIH’s OD **R24** grant OD031956 to LP, also CNPQ (313023/2020-4 and 313588/2025-2).

