## Supplementary Figures for "Whole-Embryo 3D Quantification Reveals Flexible Cellular Scaling and Conserved Tissue Architecture in *Xenopus* Species"

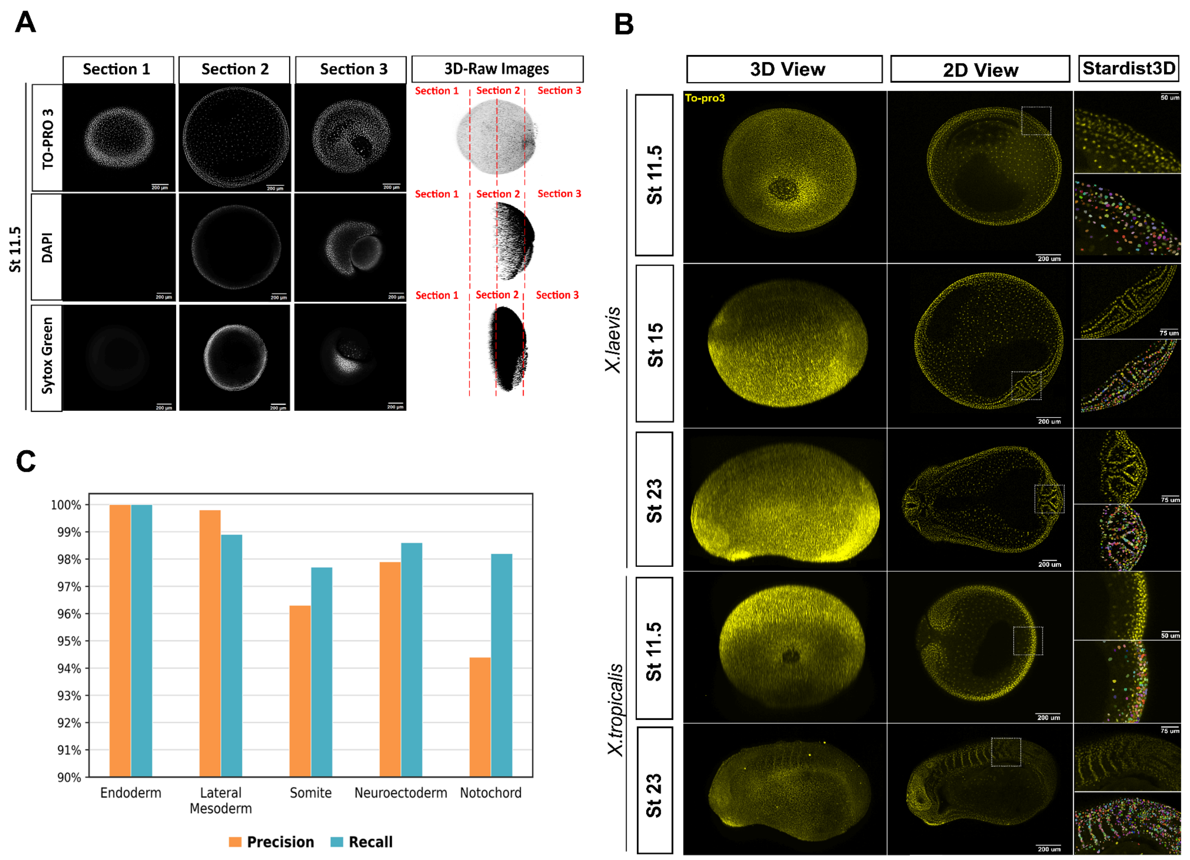


**Supplementary Figure 1.High-throughput optical clearing and 3D nuclear segmentation enable accurate cell quantification across *Xenopus* embryonic development.** (A) Comparison of nuclear dyes (TO-PRO-3, DAPI, and Sytox Green) across sequential optical sections and 3D raw reconstructions, highlighting optimal signal depth and homogeneity achieved with TO-PRO-3 staining in optically cleared embryos. (B) Representative 3D renderings, 2D mid-sagittal optical sections, and StarDist3D nuclear segmentation outputs across developmental stages (St. 11.5 to St. 23) in *Xenopus laevis* and *Xenopus tropicalis*. (C) Quantitative segmentation accuracy evaluated by Precision (orange) and Recall (blue) metrics across distinct embryonic tissue types (Endoderm, Lateral Mesoderm, Somite, Neuroectoderm, and Notochord), consistently exceeding 94% across all regions. Scale bars as indicated in individual panels.

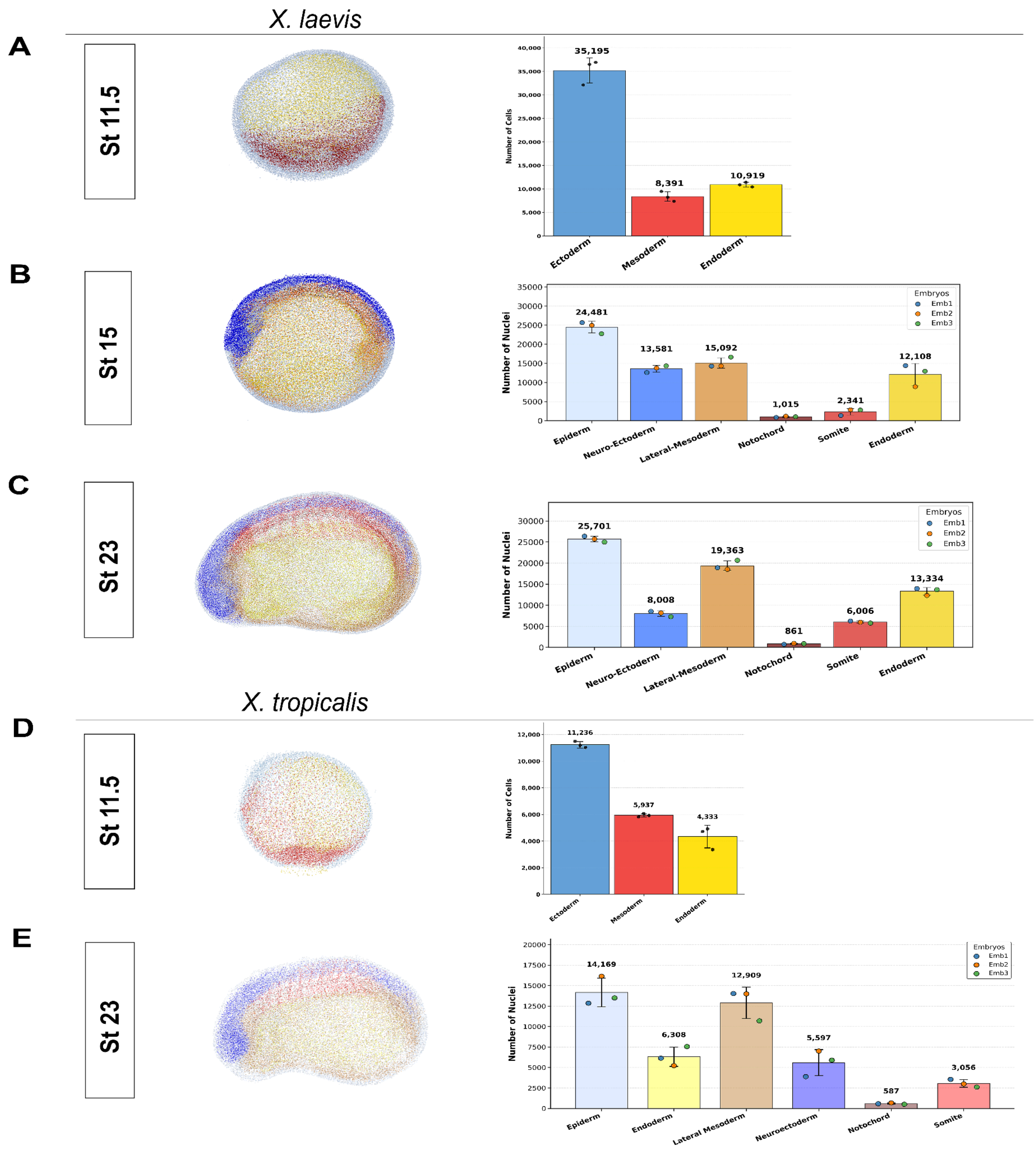


**Supplementary Figure 2. Quantitative cell number dynamics and spatial tissue allocation across developmental stages in *Xenopus laevis* and *Xenopus tropicalis*. (A–C)** Spatial nuclear distribution maps (left) alongside absolute cell counts (right) for primary germ layers and differentiated tissue domains across key developmental stages in *Xenopus laevis*: gastrula stage 11.5 (**A**), neurula stage 15 (**B**), and tailbud stage 23 (**C**). Data highlight the progressive lineage-specific cell accumulation, transitioning from initial broad germ layer compartmentalization (Ectoderm, Mesoderm, Endoderm) at St 11.5 to detailed tissue subdivisions (Epiderm, Neuro-Ectoderm, Lateral-Mesoderm, Notochord, Somite, Endoderm) at later stages. **(D–E)** Spatial nuclear point clouds and tissue-specific nuclear quantification for corresponding stages in *Xenopus tropicalis*, including stage 11.5 (**D**) and stage 23 (**E**). Bar plots represent mean nuclear counts across individual replicates (n = 3 embryos per stage, indicated by individual data points), with error bars depicting standard error of the mean (SEM).

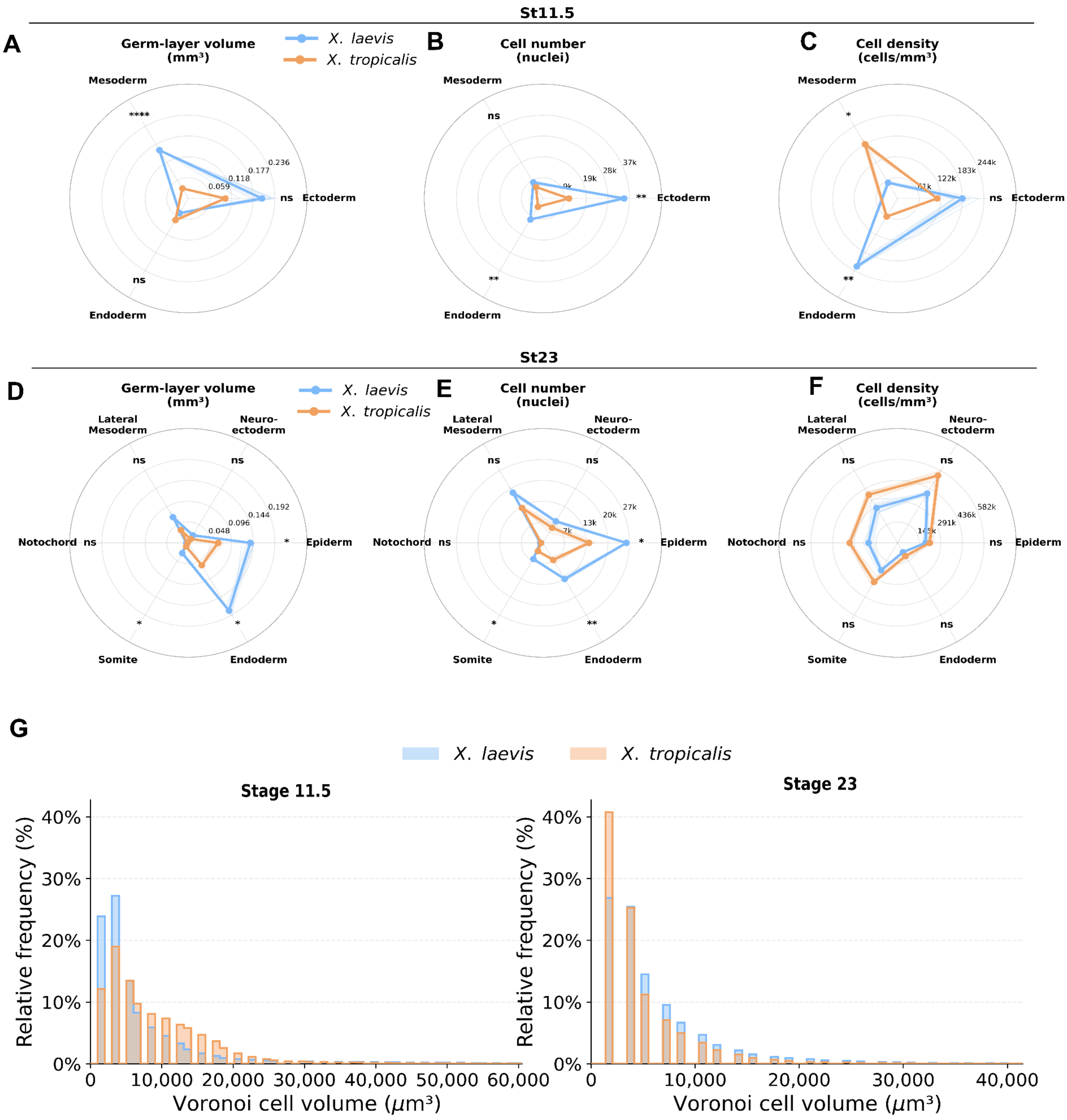


**Supplementary Figure 3. A shift in Germ-layer volume and cell size reveal a convergence between species at tailbud stage.** (A–C) Radar plots comparing overall germ-layer volume (A, $mm^3$), absolute nuclear cell number (B), and calculated cell packing density (C, cells/$mm^3$) between *X. laevis* (blue) and *X. tropicalis* (orange) across primary germ layers (Ectoderm, Mesoderm, Endoderm) at Stage 11.5 (Gastrula). (D–F) Radar plots comparing overall volume (D), cell number (E), and cell density (F) across differentiated tissue sub-lineages (Neuroectoderm, Epiderm, Endoderm, Somite, Notochord, and Lateral Mesoderm) at Stage 23 (Tailbud). (G) Relative frequency histograms (%) showing the distribution of calculated 3D Voronoi cell volumes ($\mu m^3$) for *X. laevis* (blue) and *X. tropicalis* (orange) at Stage 11.5 (left) and Stage 23 (right), highlighting the developmental shift toward smaller cell sizes and altered volume heterogeneity in both species. Statistical significance indicated on radar axes: *$p < 0.05$, $p < 0.01$, ****$p < 0.0001$, ns = not significant (mean ± SD, n = 3 embryos per stage/species).

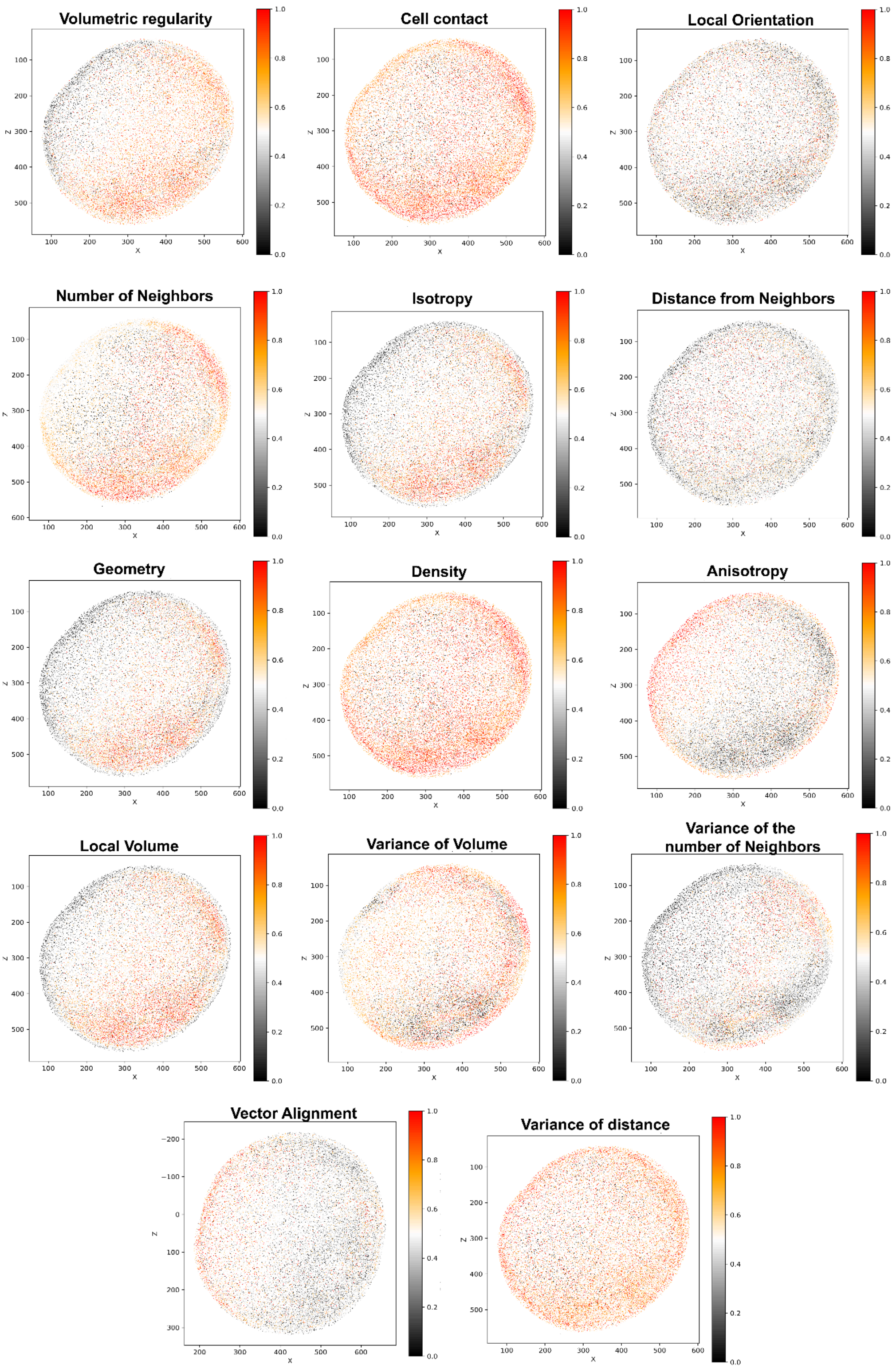


**Supplementary Figure 4. Whole-Embryo 3D Spatial Projections of 14 Local Neighborhood Morphometric Features at *X. laevis* ST 11.5.** Continuous 3D spatial mapping of the 14 individual local neighborhood morphometric parameters extracted from nuclear coordinates across the embryonic volume. Individual cellular coordinates are color-coded according to their normalized values on a dimensionless 0 to 1 scale (ranging from 0.0 in dark grey/black to 1.0 in red/orange), depicting volumetric regularity (uniformity of local cell spacing), cell contact (inverse squared inter-nuclear distance as a proxy for proximity), local orientation (spatial alignment of local cell clusters relative to embryonic axes), number of neighbors (count of neighboring nuclei within a fixed Euclidean radius), isotropy (ratio of smallest to largest eigenvalues from the 3D covariance matrix), distance from neighbors (mean Euclidean distance), geometry (local 3D cellular arrangement shape), density (local cell concentration), anisotropy (directional tissue alignment from primary eigenvalues), local volume (estimated cell volume), variance of volume (spatial variation in local cell volume), variance of the number of neighbors (fluctuations in neighbor counts), vector alignment (degree of parallel alignment among neighboring displacement vectors), and variance of distance (dispersion of inter-nuclear distances within local neighborhoods). All continuous metrics were clipped at the 5th and 95th percentiles and normalized via Min-Max rescaling (0–1 scale) to enable spatial integration and cross-sample comparison.

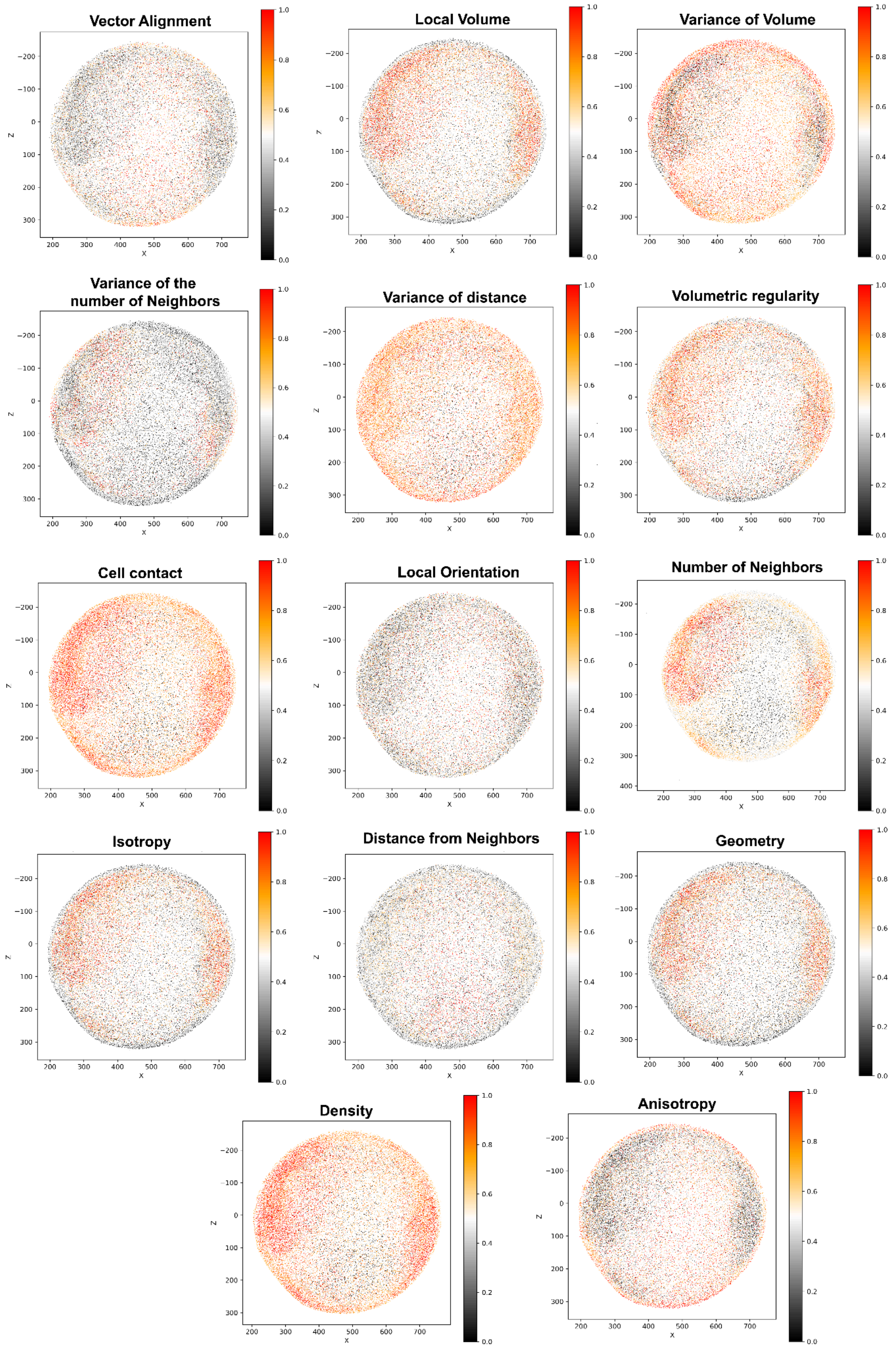

**Supplementary Figure 5. Whole-Embryo 3D Spatial Projections of 14 Local Neighborhood Morphometric Features at *X. laevis* St 15.** Continuous 3D spatial mapping of the 14 individual local neighborhood morphometric parameters extracted from nuclear coordinates across the embryonic volume. Individual cellular coordinates are color-coded according to their normalized values on a dimensionless 0 to 1 scale (ranging from 0.0 in dark grey/black to 1.0 in red/orange), depicting volumetric regularity (uniformity of local cell spacing), cell contact (inverse squared inter-nuclear distance as a proxy for proximity), local orientation (spatial alignment of local cell clusters relative to embryonic axes), number of neighbors (count of neighboring nuclei within a fixed Euclidean radius), isotropy (ratio of smallest to largest eigenvalues from the 3D covariance matrix), distance from neighbors (mean Euclidean distance), geometry (local 3D cellular arrangement shape), density (local cell concentration), anisotropy (directional tissue alignment from primary eigenvalues), local volume (estimated cell volume), variance of volume (spatial variation in local cell volume), variance of the number of neighbors (fluctuations in neighbor counts), vector alignment (degree of parallel alignment among neighboring displacement vectors), and variance of distance (dispersion of inter-nuclear distances within local neighborhoods). All continuous metrics were clipped at the 5th and 95th percentiles and normalized via Min-Max rescaling (0–1 scale) to enable spatial integration and cross-sample comparison.

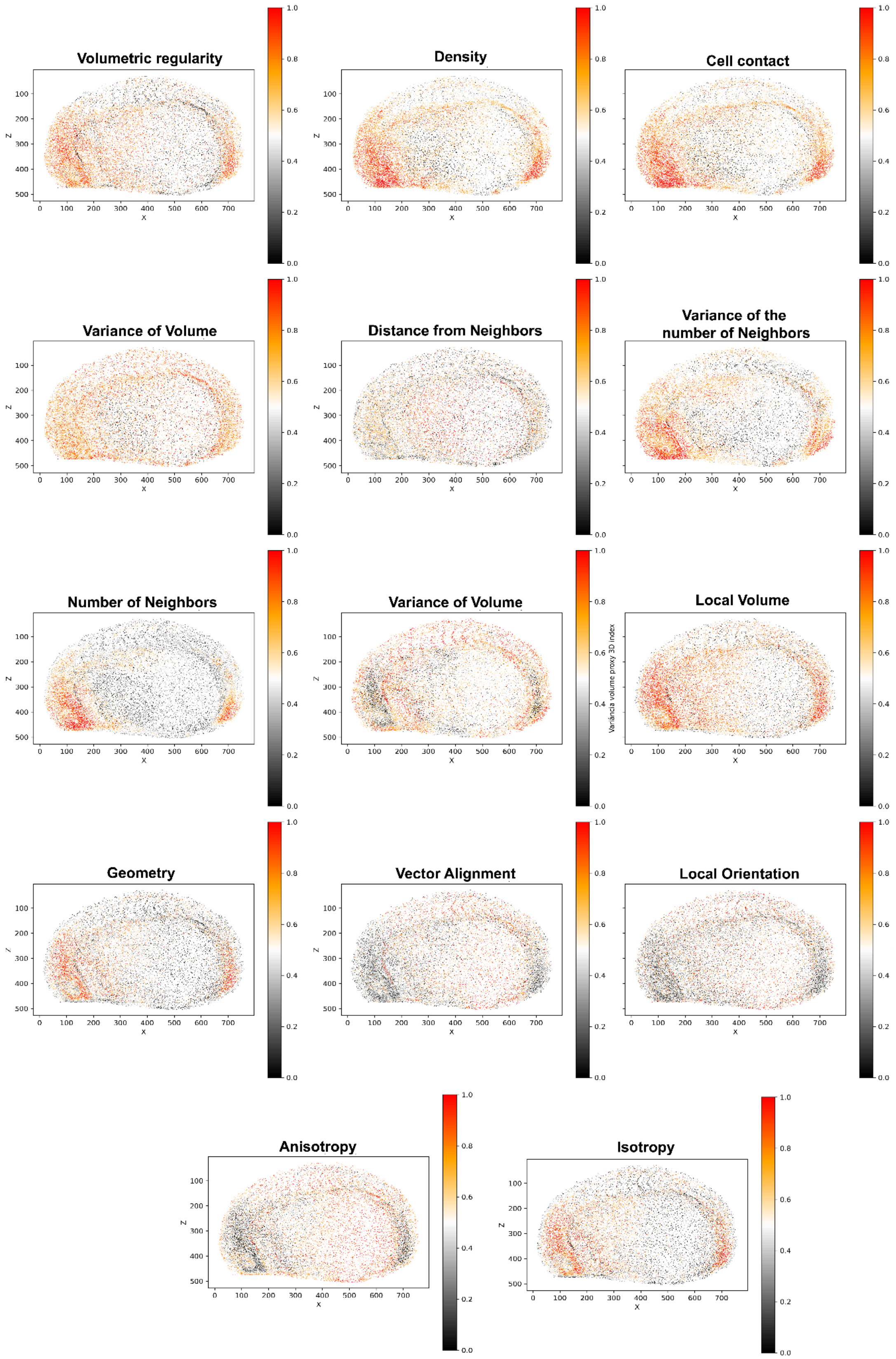


**Supplementary Figure 6. Whole-Embryo 3D Spatial Projections of 14 Local Neighborhood Morphometric Features at *X. laevis* St 23.** Continuous 3D spatial mapping of the 14 individual local neighborhood morphometric parameters extracted from nuclear coordinates across the embryonic volume. Individual cellular coordinates are color-coded according to their normalized values on a dimensionless 0 to 1 scale (ranging from 0.0 in dark grey/black to 1.0 in red/orange), depicting volumetric regularity (uniformity of local cell spacing), cell contact (inverse squared inter-nuclear distance as a proxy for proximity), local orientation (spatial alignment of local cell clusters relative to embryonic axes), number of neighbors (count of neighboring nuclei within a fixed Euclidean radius), isotropy (ratio of smallest to largest eigenvalues from the 3D covariance matrix), distance from neighbors (mean Euclidean distance), geometry (local 3D cellular arrangement shape), density (local cell concentration), anisotropy (directional tissue alignment from primary eigenvalues), local volume (estimated cell volume), variance of volume (spatial variation in local cell volume), variance of the number of neighbors (fluctuations in neighbor counts), vector alignment (degree of parallel alignment among neighboring displacement vectors), and variance of distance (dispersion of inter-nuclear distances within local neighborhoods). All continuous metrics were clipped at the 5th and 95th percentiles and normalized via Min-Max rescaling (0–1 scale) to enable spatial integration and cross-sample comparison.

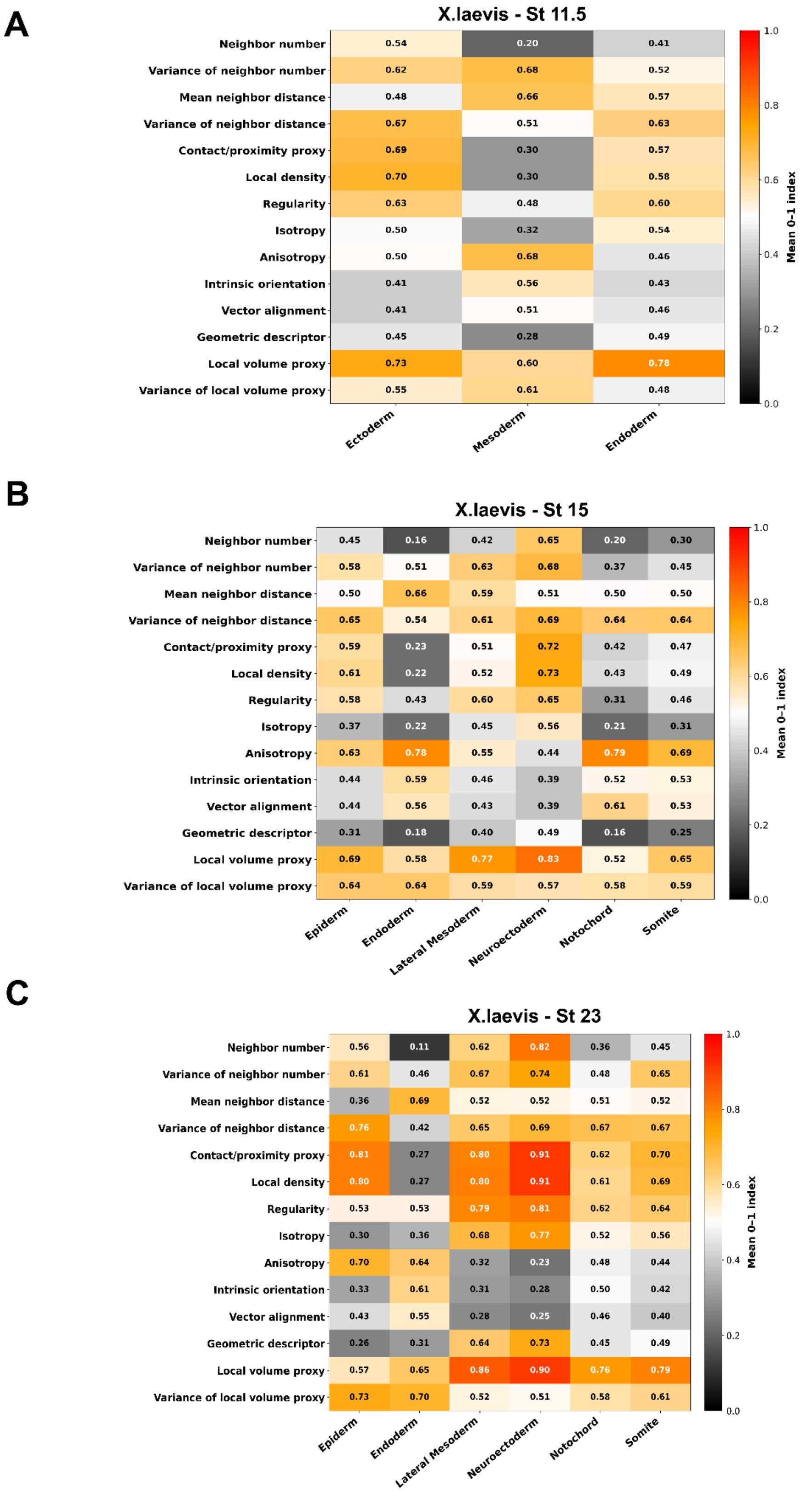


**Supplementary Figure 7. Heatmaps of Mean Morphometric Features Across Tissue Lineages in *Xenopus laevis*.** Heatmaps displaying the mean values of 14 local 3D morphometric parameters calculated across germ layers and differentiated tissue lineages during *Xenopus laevis* development. Normalized index values range from 0.0 (dark grey/black) to 1.0 (bright orange/red). **(A)** Gastrula stage (Stage 11.5) tissue-level profile across the primary germ layers (Ectoderm, Mesoderm, and Endoderm). **(B)** Neurula stage (Stage 15)

tissue-level profile across differentiated lineages (Epiderm, Endoderm, Lateral Mesoderm, Neuroectoderm, Notochord, and Somite). **(C)** Early tailbud/larval stage (Stage 23) tissue-level profile across the corresponding differentiated compartments. These quantifications track dynamic architectural shifts in local neighborhood properties, such as cell packing, volume, and alignment, across developmental stages and tissue lineages.

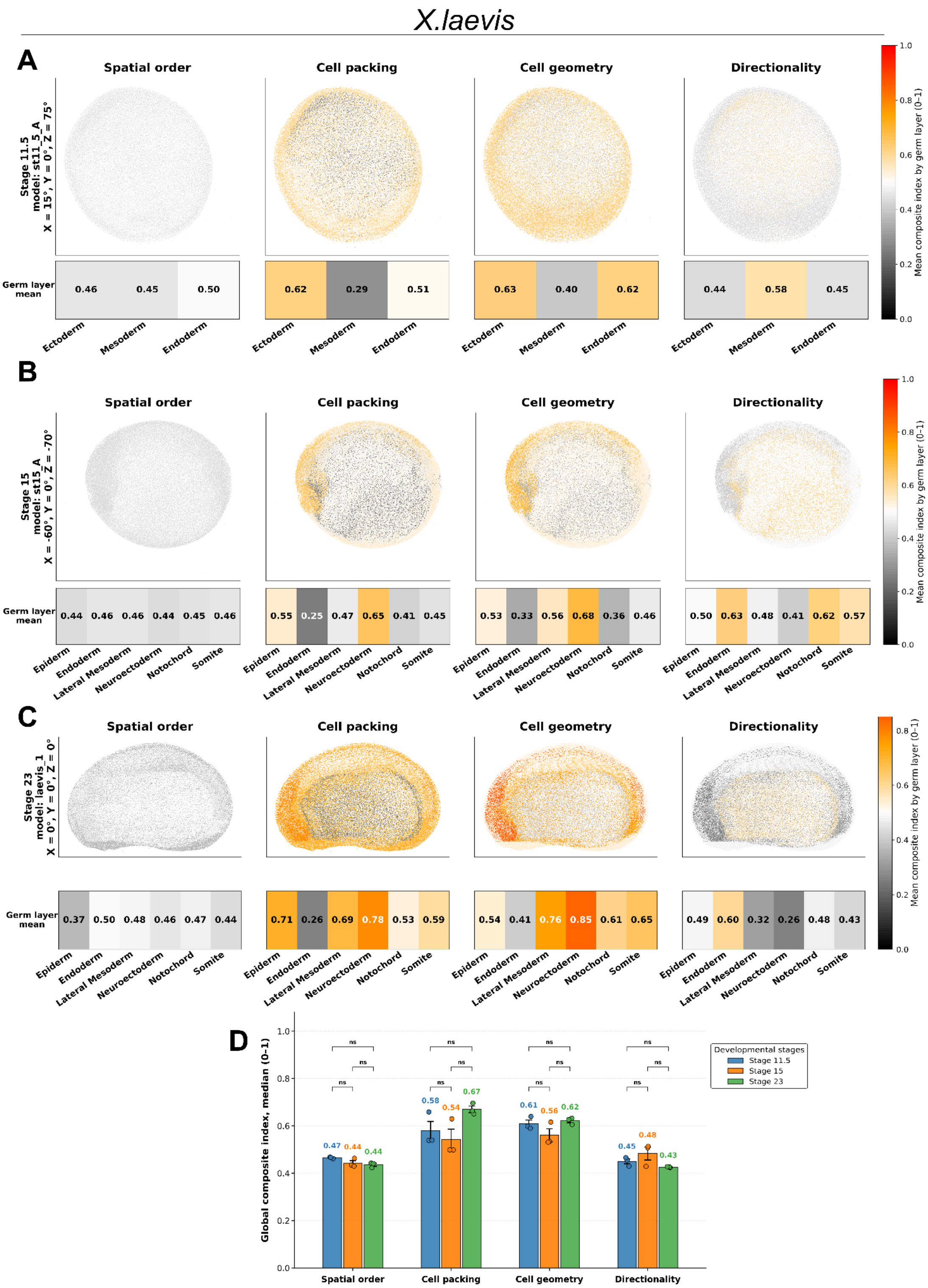


**Supplementary Figure 8. Spatiotemporal cellular architecture mapping reveals shifting physical and architectural dynamics during *Xenopus laevis* development. (A–C)** Spatial projection and tissue quantification of composite organizational indices. The 14 local neighborhood morphometric features are integrated into four functional macro-categories (*Spatial order*, *Cell packing*, *Cell geometry*, and *Directionality*). These composite indices are projected as continuous spatial gradients onto 3D embryo coordinates (top) and quantified as mean values across germ layers/tissues (bottom) for gastrulation at St. 11.5 **(A)**, neurulation at St.

15 **(B)**, and larval stage at St. 23 **(C)** (n = 3 per stage). **(D)** Global architectural dynamics. Multi-parametric comparison of global composite indices (median values 0–1) across the three developmental stages. Bars represent the mean ± SEM, with individual data points corresponding to single embryos. Pairwise statistical comparisons were conducted using two-sided Welch's *t*-tests with Benjamini-Hochberg false discovery rate (FDR-BH) adjustment across 12 tests (* $q < 0.05$; $q < 0.01$; ns, not significant). This global quantification tracks macro-scale shifts in physical tissue cellular architecture, highlighting the progressive structural reorganization of the embryo over time.

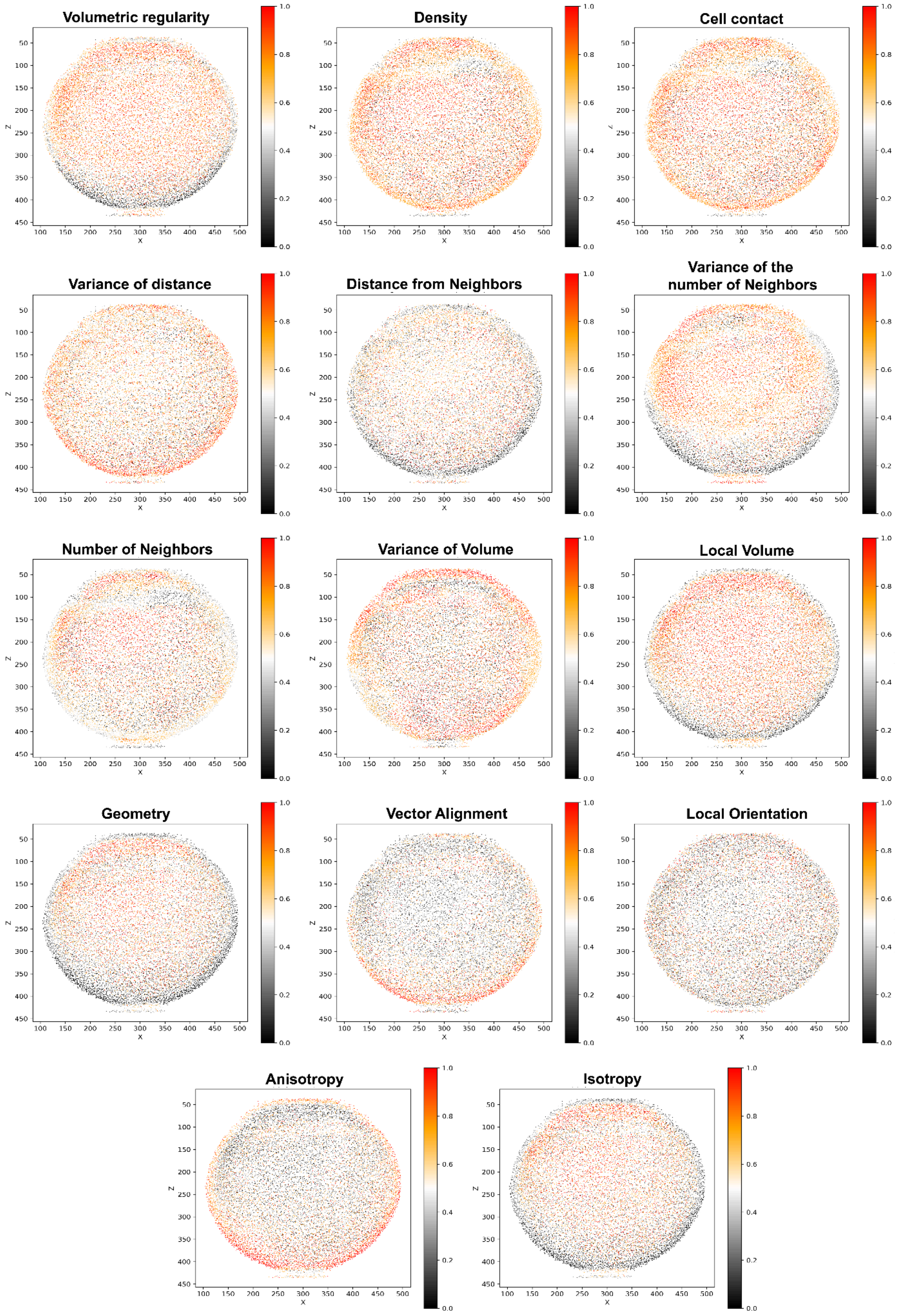

**Supplementary Figure 9. Whole-Embryo 3D Spatial Projections of 14 Local Neighborhood Morphometric Features at *X. tropicalis* ST 11.5.** Continuous 3D spatial mapping of the 14 individual local neighborhood morphometric parameters extracted from nuclear coordinates across the embryonic volume. Individual cellular coordinates are color-coded according to their normalized values on a dimensionless 0 to 1 scale (ranging from 0.0 in dark grey/black to 1.0 in red/orange), depicting volumetric regularity (uniformity of local cell spacing), cell contact (inverse squared inter-nuclear distance as a proxy for proximity), local orientation (spatial alignment of local cell clusters relative to embryonic axes), number of neighbors (count of neighboring nuclei within a fixed Euclidean radius), isotropy (ratio of smallest to largest eigenvalues from the 3D covariance matrix), distance from neighbors (mean Euclidean distance), geometry (local 3D cellular arrangement shape), density (local cell concentration), anisotropy (directional tissue alignment from primary eigenvalues), local volume (estimated cell volume), variance of volume (spatial variation in local cell volume), variance of the number of neighbors (fluctuations in neighbor counts), vector alignment (degree of parallel alignment among neighboring displacement vectors), and variance of distance (dispersion of inter-nuclear distances within local neighborhoods). All continuous metrics were clipped at the 5th and 95th percentiles and normalized via Min-Max rescaling (0–1 scale) to enable spatial integration and cross-sample comparison.

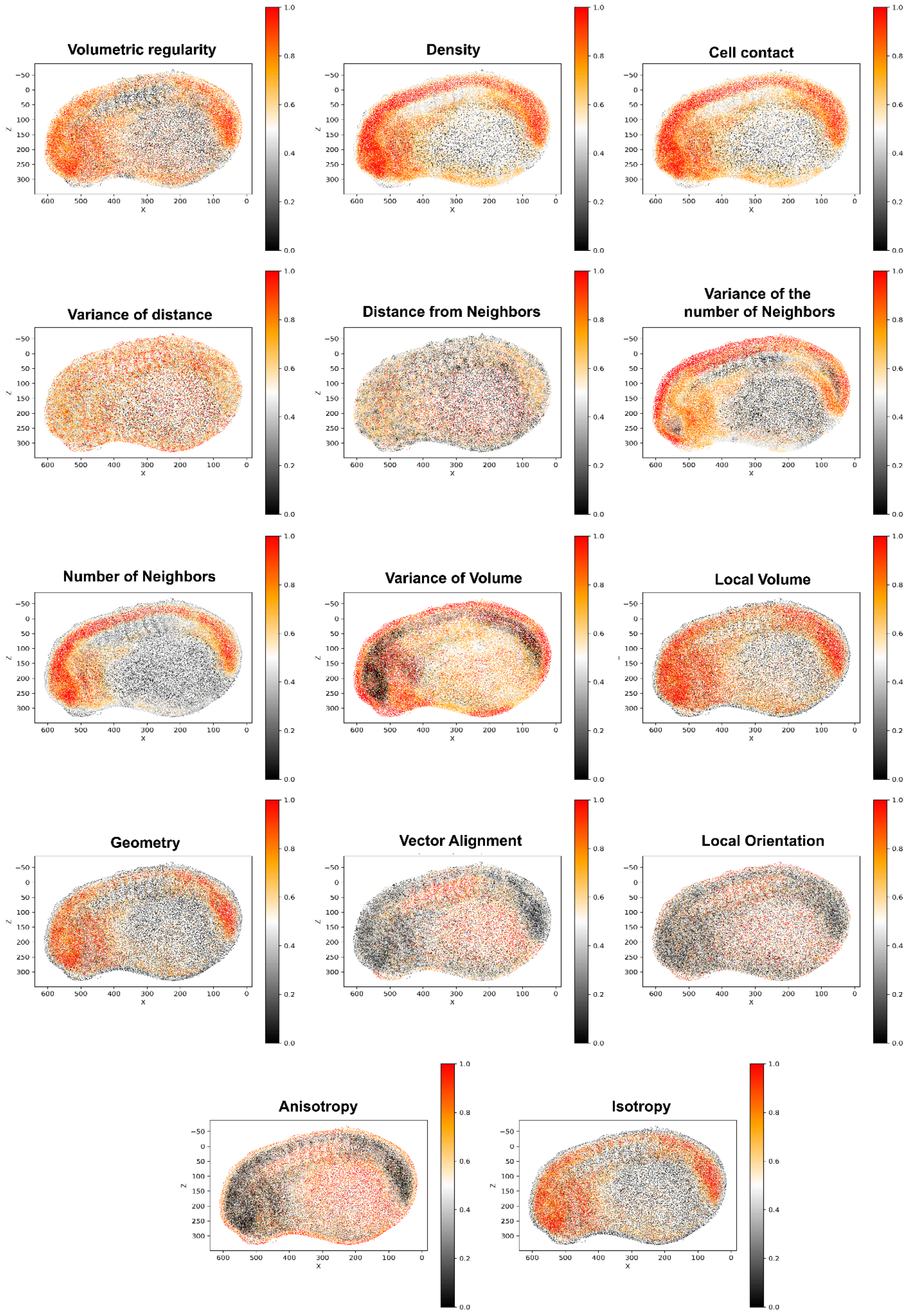


**Supplementary Figure 10. Whole-Embryo 3D Spatial Projections of 14 Local Neighborhood Morphometric Features at *X. tropicalis* St 23.** Continuous 3D spatial mapping of the 14 individual local neighborhood morphometric parameters extracted from nuclear coordinates across the embryonic volume. Individual cellular coordinates are color-coded according to their normalized values on a dimensionless 0 to 1 scale (ranging from 0.0 in dark grey/black to 1.0 in red/orange), depicting volumetric regularity (uniformity of local cell spacing), cell contact (inverse squared inter-nuclear distance as a proxy for proximity), local orientation (spatial alignment of local cell clusters relative to embryonic axes), number of neighbors (count of neighboring nuclei within a fixed Euclidean radius), isotropy (ratio of smallest to largest eigenvalues from the 3D covariance matrix), distance from neighbors (mean Euclidean distance), geometry (local 3D cellular arrangement shape), density (local cell concentration), anisotropy (directional tissue alignment from primary eigenvalues), local volume (estimated cell volume), variance of volume (spatial variation in local cell volume), variance of the number of neighbors (fluctuations in neighbor counts), vector alignment (degree of parallel alignment among neighboring displacement vectors), and variance of distance (dispersion of inter-nuclear distances within local neighborhoods). All continuous metrics were clipped at the 5th and 95th percentiles and normalized via Min-Max rescaling (0–1 scale) to enable spatial integration and cross-sample comparison.

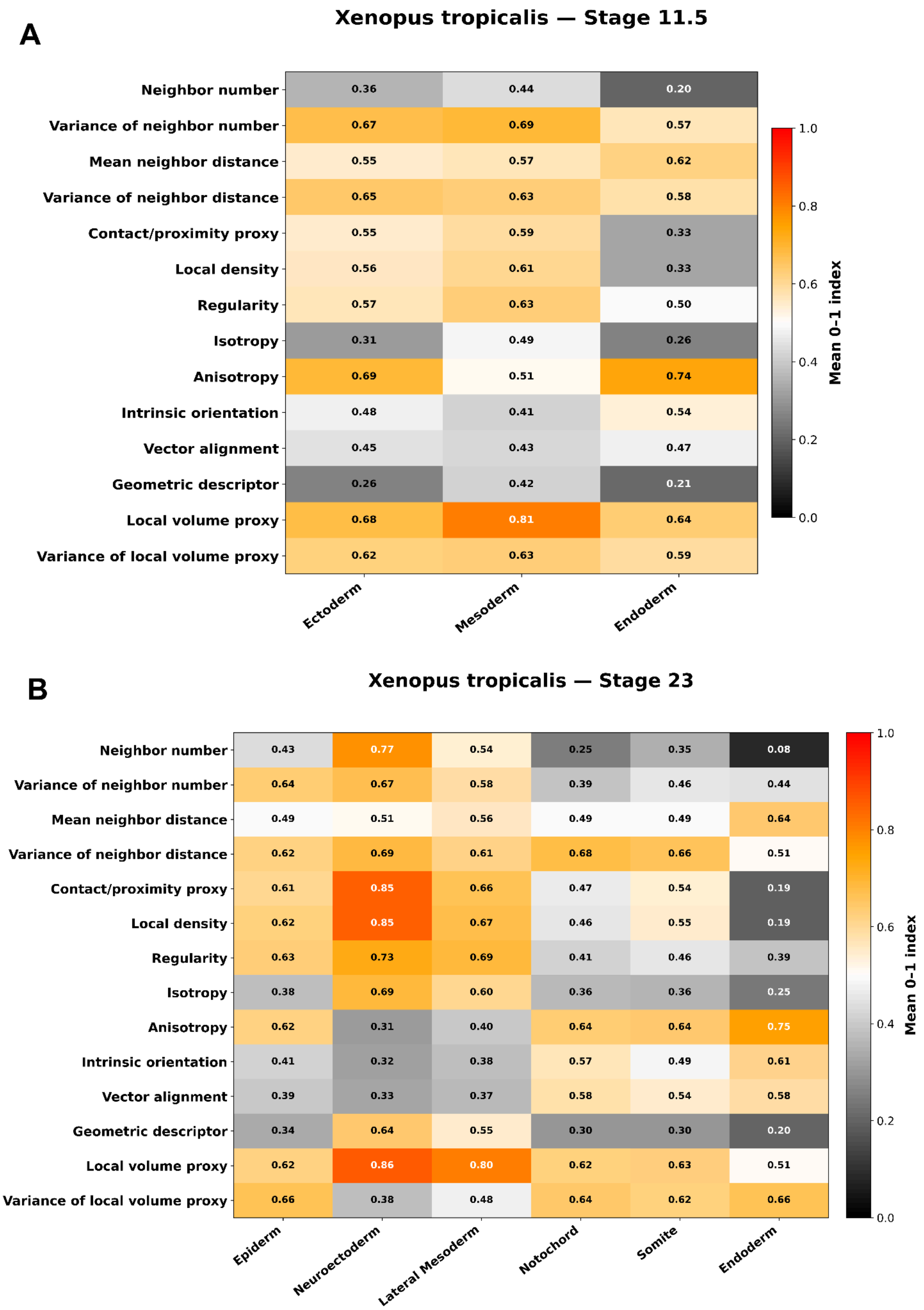


**Supplementary Figure 11. Heatmaps of Mean Morphometric Features Across Tissue Lineages in *Xenopus tropicalis*.** Heatmaps displaying the mean values of 14 local 3D morphometric parameters calculated across germ layers and differentiated tissue lineages during *Xenopus tropicalis* development. Normalized index values range from 0.0 (dark grey/black) to 1.0 (bright orange/red). **(A)** Gastrula stage (Stage 11.5) tissue-level profile across the primary germ layers (Ectoderm, Mesoderm, and Endoderm). **(B)** Early tailbud/larval stage (Stage 23) tissue-level profile across differentiated compartments (Epiderm, Neuroectoderm, Lateral Mesoderm, Notochord, Somite, and Endoderm). These quantifications provide a lineage-specific topological baseline for *Xenopus tropicalis*, highlighting architectural similarities and conserved tissue organization relative to *Xenopus laevis*.

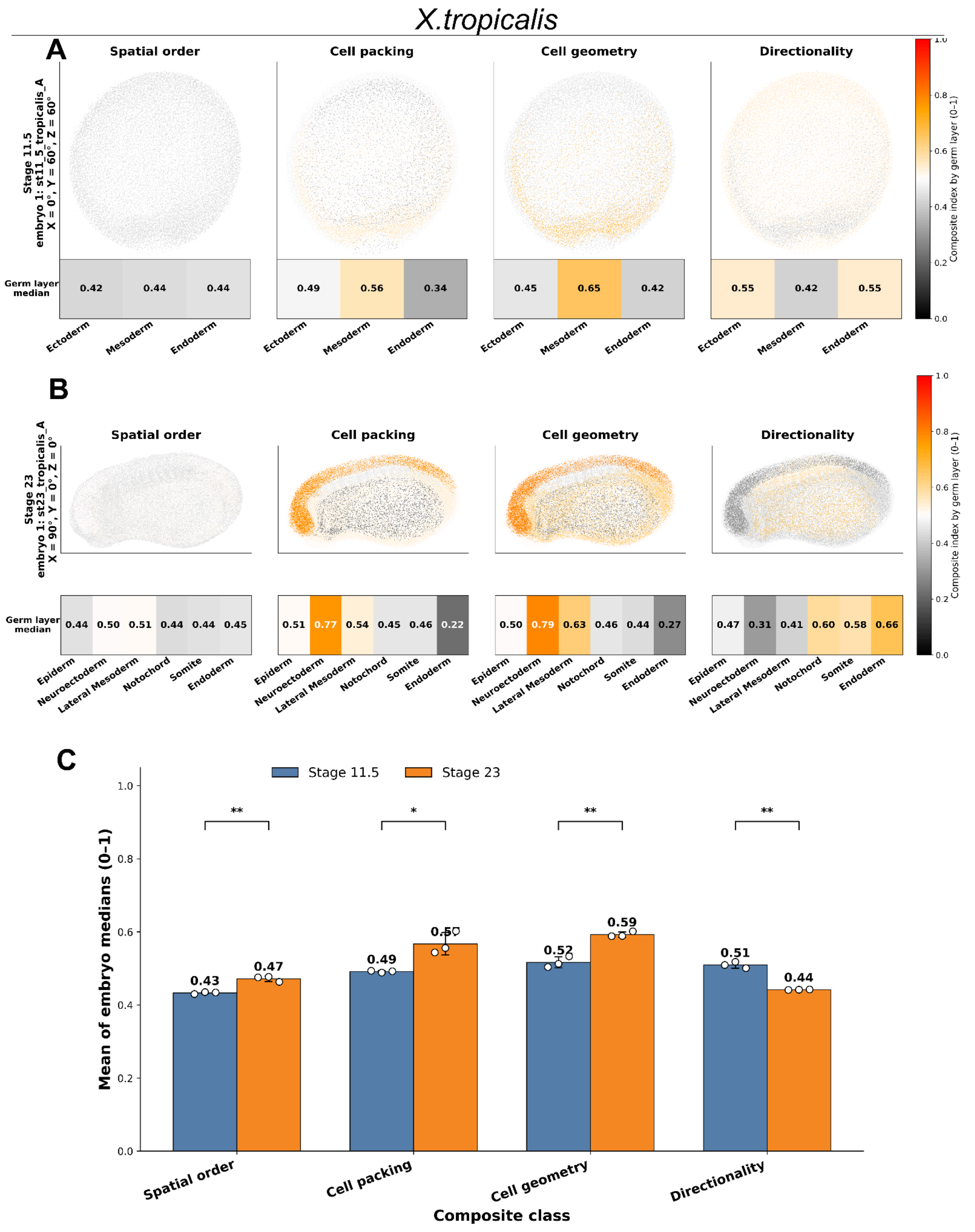


**Supplementary Figure 12. Quantitative architectural profiling and developmental progression of tissue organization parameters in *Xenopus tropicalis*. (A)** Spatial distribution maps and germ layer median values for key morphological feature classes—Spatial order, Cell packing, Cell geometry, and Directionality—in representative *Xenopus tropicalis* embryos at stage 11.5 across Ectoderm, Mesoderm, and Endoderm. **(B)** Corresponding spatial maps and tissue-level median scores for stage 23 embryos across expanded tissue domains (Epiderm, Neuroectoderm, Lateral Mesoderm, Notochord, Somite, and Endoderm), displaying localized architectural refinement. **(C)** Quantitative comparison of whole-embryo median composite indices between stage 11.5 (blue) and stage 23 (orange), showing significant increases in Spatial order, Cell packing, and Cell geometry alongside a decrease in Directionality (*p < 0.05, p < 0.01).

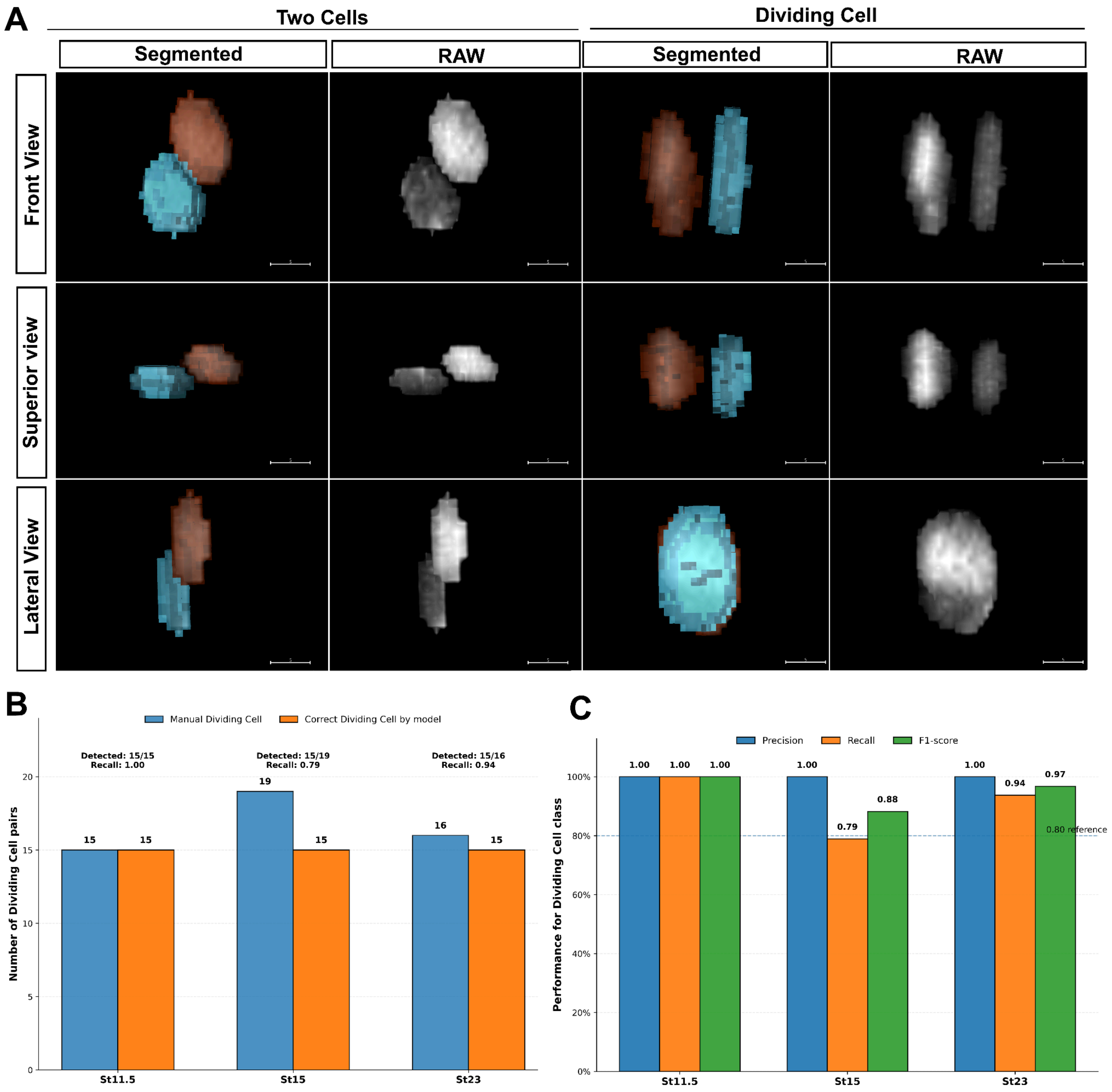


**Supplementary Figure 13. Morphological Characterization and Model Performance Validation for Automated Mitotic Cell Classification. (A)** Comparative 3D orthogonal views (Front, Superior, and Lateral views) of segmented masks and corresponding raw nuclear fluorescence (RAW) contrasting adjacent non-dividing pairs ("Two Cells") against actively dividing sister nuclei ("Dividing Cell"). Segmented panels display distinct colored 3D masks (orange and cyan) highlighting the spatial orientation and geometry characteristic of mitotic pairs. **(B)** Quantitative benchmark comparing manual expert annotations of dividing cell pairs (blue) against correct classifications identified by the automated computational model (orange) across developmental stages St. 11.5, St. 15, and St. 23. Detected pair ratios and corresponding recall rates are indicated for each stage. **(C)** Classification performance metrics, Precision (blue), Recall (orange), and F1-score (green), for the dividing cell class across developmental stages, demonstrating high precision (1.00 across all stages) and robust F1-scores exceeding the 0.85 performance reference baseline.

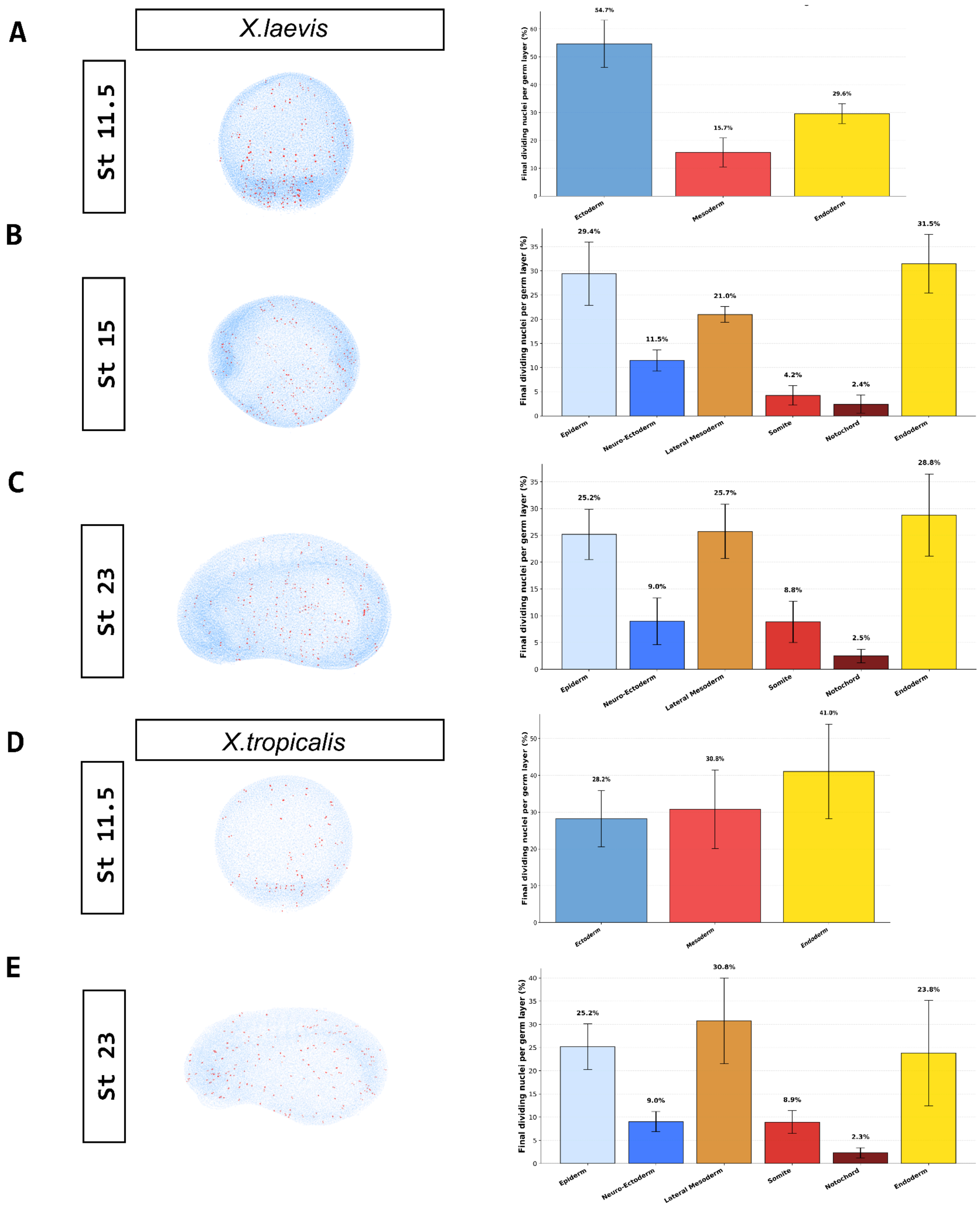


**Supplementary Figure 14. Comparative spatiotemporal distribution and germ layer-specific proportions of mitotic events during *Xenopus* morphogenesis. (A–C)** Spatial mapping of dividing nuclei (left) and corresponding percentage contributions per germ layer/tissue domain (right) in *Xenopus laevis* across gastrula stage 11.5 (**A**), neurula stage 15 (**B**), and tailbud stage 23 (**C**). Data depict a dynamic reallocation of active cell divisions from early ectodermal dominance at stage 11.5 toward prominent endodermal and mesodermal fractions in later stages. **(D–E)** Spatial distribution maps and quantitative tissue allocation of mitotic nuclei in *Xenopus tropicalis* at stage 11.5 (**D**) and stage 23 (**E**), highlighting conserved cell division patterns between species across equivalent developmental windows. Bar plots represent mean percentages of total dividing nuclei per region, with error bars indicating SEM.

**Table. Feature domains used to classify dividing-cell candidates**

| Feature domain | Representative variables | Criterion supporting Dividing Cell |
|---|---|---|
| **Pair proximity** | pair_distance_norm; contact_distance_norm; surface_distance_norm; bbox_gap_norm; dilated_contact_fraction | The two labels are close, with limited gap and plausible physical contact. |
| **Size similarity** | volume_similarity; fast_volume_similarity; internal_axis_length_similarity | The two nuclei have comparable volume and internal-axis length. |
| **Shape and axis similarity** | sphericity_similarity; elongation_similarity; axis_similarity; internal_axis_similarity; v11_axis_between_score | The nuclei show compatible shape and major-axis orientation. |
| **Intensity similarity** | mean_intensity_similarity; max_intensity_similarity; intensity_histogram_similarity; intensity_p90_similarity | The two nuclei have similar fluorescence intensity profiles. |
| **Mirror symmetry** | mirror_shape_similarity; mirror_intensity_similarity; mirror_intensity_correlation01; mirror_symmetry_score | Mirroring one nucleus across the pair midpoint matches the other in shape and intensity. |
| **Three-level geometry** | three_level_non_crossing_alignment_score; three_level_internal_axis_alignment_score; three_level_distance_cv_score; three_level_parallelism_score; n_crossings_with_middle | Top, middle and bottom levels align in an ordered, parallel and non-crossing configuration. |
| **Interface intensity** | bridge_mean_ratio; bridge_min_ratio; line_middle_mean_ratio; bridge_low_score | The intensity profile between nuclei is compatible with a division interface. |
| **Composite sister scores** | sister_consensus_score; mirror_consensus_score; three_level_consensus_score; high_confidence_sister_score | Integrated biological scores support a sister-nucleus signature. |
| **Manual/prototype scores** | manual_supervised_score; positive_prototype_score; negative_prototype_score; discriminative_prototype_score; prototype_margin_score; final_division_quality_score | The pair resembles manually annotated Dividing Cell examples more than Two cells. |
| **Final decision** | probability_dividing_cell plus selected biological gate features | A pair is accepted when model probability and biological consistency are both high. |

**Supplementary Table 1. Feature Groups and Biological Rationales Used for Automated Classification of Dividing-Cell Candidates.** The V12B classifier employs ten interpretable feature groups to distinguish sister-nucleus dividing pairs from non-dividing pairs. These comprise pair proximity and contact metrics for spatial closeness, size and volume similarity for comparable nuclear dimensions, and shape and axis similarity for geometric compatibility. Additional parameters evaluate intensity similarity across fluorescence profiles, mirror symmetry for shape and intensity overlap, and three-level internal-axis geometry for ordered non-crossing alignment. Finally, the model uses bridge and interface intensity profiles for division lines, integrates composite sister-nucleus scores, incorporates manual/prototype-derived scores from expert annotations, and applies a final classification rule combining model probability with biological gates. Stage and sample identities are excluded from biological features to ensure classification neutrality
